# PHbinder and PSGM: A Cascaded Framework for Epitope Prediction and HLA-I Allele Identification

**DOI:** 10.1101/2025.06.25.661428

**Authors:** Zikun Wang, Xueying Wang, Jixiu Zhai, Shengrui Xu, Tianchi Lu

## Abstract

The presentation of antigens by Human Leukocyte Antigen class I (HLA-I) molecules is a cornerstone of adaptive immunity. Although existing prediction tools such as NetMHCpan and MHCflurry exhibit high accuracy in predicting binding affinity between peptides and specific HLA-I alleles, they are constrained to a preset set of alleles. Consequently, they can neither directly determine whether a peptide is an epitope nor provide a holistic binding profile across the entire HLA-I allelic landscape. To overcome these challenges, we introduce two synergistic models: PHbinder (Peptide-HLA-I Binder) and PSGM (Pseudo Sequence Generation and Mapping). PHbinder integrates features from a fine-tuned ESM2 language model with Low-Rank Adaptation (LoRA), processing them through parallel CNN and Transformer branches to capture local and global patterns, which are then fused using a Cross-Multi-Head Attention mechanism. In the epitope prediction task, PHbinder achieved an accuracy of prediction of 85. 12%, significantly exceeding established benchmark models. Complementing this, the PSGM model employs a Generative Adversarial Network(GAN) architecture to generate the corresponding HLA-I pseudo sequences. These are then mapped to the known alleles using a Hamming distance-based nearestneighbor search. PSGM achieved 49.26% average coverage in its predictions of the Top-50 alleles. Furthermore, orthogonal validation with MHCflurry revealed that 63% of the highest affinity binding partners within its Top-50 list were new experimentally unverified HLA-I alleles. Together, PHbinder and PSGM establish a cascaded framework that enables a precise “Peptide → Epitope Determination → HLA-I Alleles List” pipeline. This work accelerates the screening of immunogenic epitopes and provides a powerful upstream preprocessor for traditional prediction tools.

## Introduction

Human Leukocyte Antigen class I (HLA-I)[1] molecules are responsible for presenting endogenous or exogenous antigenic epitopes[2] on the cell surface. This presentation allows T cells to recognize them via their specific T-cell receptors (TCR)[3][4], which in turn initiates or suppresses an immune response. This molecular recognition, centered on the complex of peptide-HLA-I (p-HLA-I), is critical for immune surveillance[5] against virus-infected cells[6] and malignant tumors[7]. Consequently, the accurate prediction of p-HLA-I binding is a foundational step for identifying immunotherapy targets, screening for antigenic epitopes, and designing next-generation vaccines[8].

In the field of prediction of p-HLA-I binding[9], early computational models date back to Parker’s Position-Specific Scoring Matrix (PSSM)[10] in 1994, which inspired methods such as BIMAS[11] and SYFPEITHI[12] that achieved respectable performance for specific alleles. The introduction of NetMHCpan[13] in 2007 marked a significant advance, enabling the first “pan-allele” predictions across a wide range of HLA-I alleles and greatly expanding the breadth of computational analysis. Subsequently, tools such as MHCflurry[14] and ImmuneApp[15] have continued to refine and enhance predictive accuracy.

However, these state-of-the-art tools present significant limitations. NetMHCpan, MHCflurry, and their contemporaries operate on a “verification-style” workflow, requiring the user to input both a peptide and a specific HLA-I allele for evaluation. To determine if a new peptide is a broad-spectrum epitope, a researcher must iteratively test it against thousands of known HLA-I alleles— A process that is computationally intensive and poorly suited for exploratory discovery of novel p-HLA-I interactions. In 2025, TransHLA[16] proposed a new paradigm: by using only the peptide as input, it enabled the direct classification of epitopes, significantly accelerating the screening process. However, the TransHLA model suffers from two key flaws. First, it employs simple feature concatenation, a naive fusion method that fails to effectively align and correlate heterogeneous features from its CNN and Transformer[17] branches. This may prevent the model from learning deeper relationships and can introduce redundancy that degrades performance. Secondly, TransHLA’s task is confined to predicting if a peptide can be presented; it cannot identify which specific HLA-I alleles are responsible for, failing to establish a direct predictive link from peptide to allele.

To address these gaps, we developed a two-part framework composed of the PHbinder and PSGM models. PHbinder achieves state-of-the-art performance in determining whether a peptide is an HLA-I-presented epitope, while PSGM establishes a direct mapping from a given peptide to its likely HLA-I binding partners.

PHbinder retains a parallel Transformer and CNN architecture but improves upon previous designs. Instead of simple concatenation, we implement a Cross-Multi-Head Attention[18] mechanism to facilitate deep, non-linear interactions between the two feature branches for a more robust fusion. To enhance PHbinder’s feature extraction capabilities, we adopted a two-stage fine-tuning strategy. We first conduct a dedicated pre-training phase, using Low-Rank Adaptation (LoRA)[19] to efficiently adapt the general ESM2 protein language model[20] to the specific domain of HLA-I presentation. PHbinder is then fine-tuned upon this specialized model, again using LoRA. Experimental results confirm this strategy not only improves computational efficiency but also boosts model performance. When compared against both advanced sequence classification models and leading p-HLAI binding prediction software, PHbinder achieves state-of-the-art (SOTA) performance in overall epitope prediction.

The PSGM model is built on a Generative Adversarial Network(GAN)[21] architecture. It uses HLA-I pseudo sequences—the 34 key amino acid residues in the HLA molecule that directly interact with the peptide as its generation target. Peptide’s information guides the generator’s autoregressive process. Our results show that PSGM exhibits strong performance across three distinct metrics: Sequence Quality, Distributional Similarity, and Novelty. Finally, we employ a Hamming[22] distance-based nearest-neighbor[23] search to efficiently map the generated pseudo sequences back to known HLA alleles, producing a ranked list of likely binders. A comprehensive evaluation shows that PSGM achieves average coverage rates of 21.72% 30.94%, and 49.26% for its Top-20, Top-30, and Top-50 predictions, respectively. Moreover, orthogonal validation with MHCflurry[24] revealed that PSGM could identify novel, experimentally unverified binding partners, which constituted 9%, 22%, 26%, and 63% of the highest-affinity alleles in the Top-5, Top-10, Top-20, and Top-50 lists, respectively.

Overall, by cascading PHbinder and PSGM, we have constructed a framework that, for the first time, actualizes a precise “Peptide → Epitope Determination → HLA-I Alleles List” prediction pipeline[25]. This transforms the study of p-HLA-I binding from a traditional “verification problem” into an efficient “discovery problem,” redefining the research paradigm. Our approach not only accelerates the screening of broad-spectrum vaccine candidates and deepens the analysis of immune escape mechanisms but also serves as a powerful upstream preprocessor for traditional tools, enhancing experimental validation efficiency and enabling the discovery of novel, high-affinity p-HLA-I binding pairs[26].

## Results

### The Workflow and Framework of PHbinder and PSGM Models

Figure 1 provides a schematic overview of the proposed computational framework, detailing the architectures of the PHbinder and PSGM models. The workflow begins with the curation and validation of experimental data from immune epitope databases such as IEDB[27] (Figure 1a). The subsequent panels (Figures1b-1e) illustrate the architecture of the complete cascaded framework.The primary objective of PHbinder is to function as a high-precision classifier, determining whether a given input peptide is an immunogenic epitope capable of being presented by HLA-I molecules. For epitopes that are positively identified by PHbinder, the PSGM model is then employed. PSGM’s objective is to generate the corresponding HLA-I pseudo- = sequences for these epitopes and subsequently map them to known HLA-I alleles, thereby predicting the specific alleles to which the peptide can bind.

**Figure 1.**
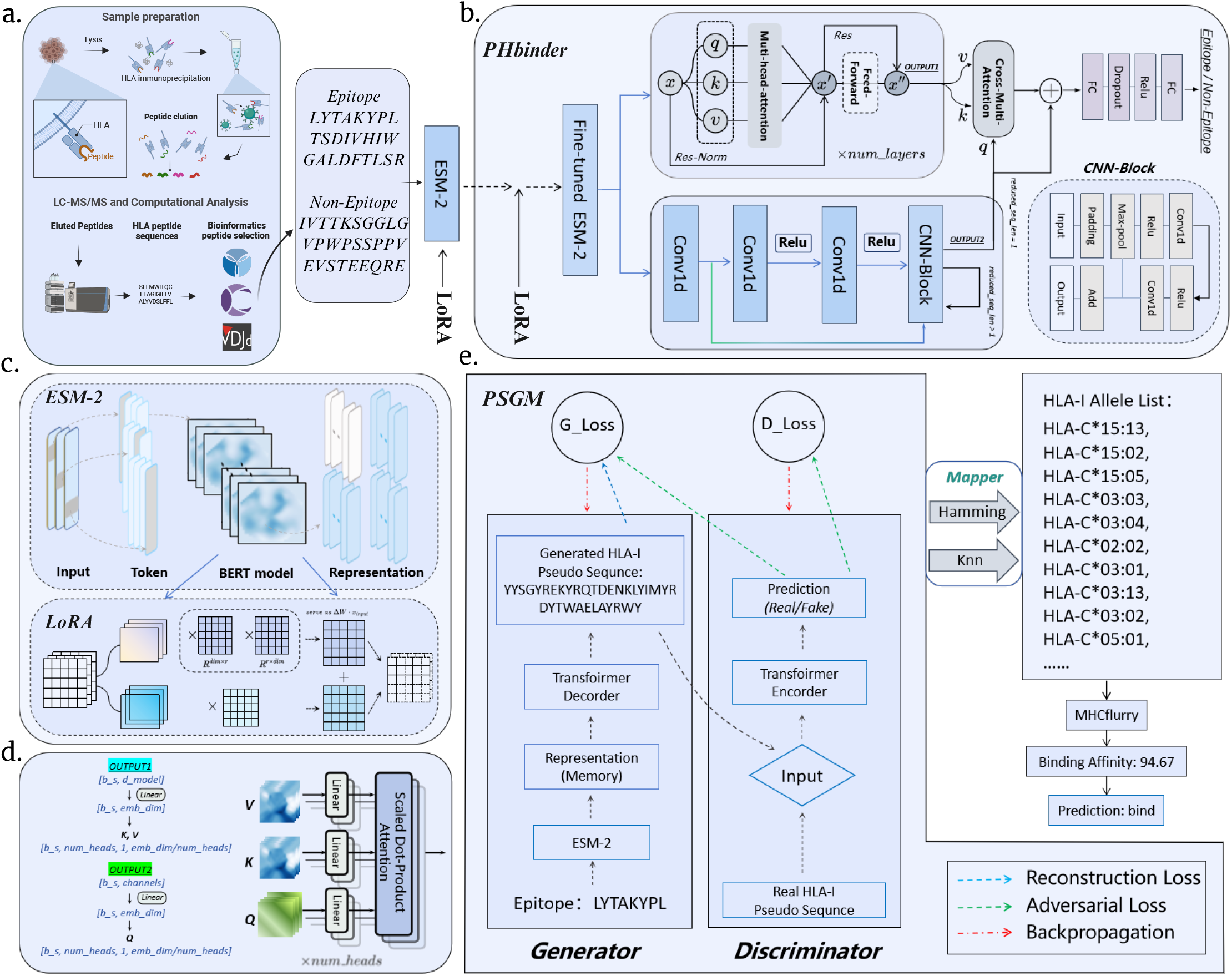
Overview of the workflow and framework for PHbinder and PSGM.a) Schematic diagram of the peptide-HLA-I binding mechanism and experimental validation data. b) The architecture of PHbinder. PHbinder takes a peptide as input, loads a pre-trained LoRA adapter into the ESM2 model, and extracts feature embeddings from the 30th layer. These embeddings are fed into parallel CNN and Transformer branches for local and global feature extraction, respectively. The resulting feature vectors are then fused using Cross-Multi-Head Attention: the local feature vector from the CNN serves as the Query (Q), while the global context vector from the Transformer serves as the Key (K) and Value (V). The resulting attention-weighted vector is concatenated with the original CNN feature vector and passed to a linear classifier to determine if the peptide is an HLA-I presented epitope. c) Detailed workflow of fine-tuning ESM2 using Low-Rank Adaptation (LoRA). d) Detailed illustration of the Cross-Multi-Head Attention mechanism. e) The architecture of PSGM. PSGM takes epitopes identified by PHbinder as input and autoregressively generates an HLA-I pseudo sequence, which is then mapped to an HLA-I Allele List. PSGM employs a GAN (Generative Adversarial Network) architecture for generation, where the Generator and Discriminator are optimized through adversarial training. The generator consists of an encoder and a 6-layer Transformer decoder. Peptide sequence features are encoded into a conditional vector, which is fed into the decoder to guide the generation process.

The construction of PHbinder is a two-stage process. First, we conduct a pre-training phase where a lightweight model, consisting of an ESM2-LoRA[20][19] module and a linear classifier, is fine-tuned on our peptide dataset. Second, the core PHbinder architecture is built upon this pre-trained ESM2-LoRA foundation. The PHbinder model extracts deep feature embeddings from the 30th layer of the ESM2 module and processes them through two parallel branches: a 1D convolutional neural network (CNN) branch to capture local motifs and a Transformer branch (a stack of six encoder layers) to capture global dependencies. To fuse these heterogeneous features, we implement a Cross-Multi-Head Attention mechanism. The local feature vector from the CNN branch serves as the Query, while the global context vector from the Transformer branch serves as both the Key and Value. The resulting attention-weighted vector, which has been dynamically refined with global context, is concatenated with the original local feature vector and passed through a feed-forward network for final classification. The detailed hyperparameter settings of PHbinder during the training process are presented in Supplementary Table S4.

The PSGM model comprises two main components: a generator for HLA-I pseudo sequence generation and a mapping module. Generation is based on a Generative Adversarial Network[21] architecture, composed of a generator and a discriminator. The generator is conditioned on an antigenic epitope; its sequence is first encoded by a pre-trained ESM2 model. This high-dimensional conditional vector, representing the peptide’s biochemical properties, is injected into a 6-layer Transformer decoder, which then autoregressively[28] generates a complete HLA-I pseudo sequence. To ensure the biological realism of the output, a 3-layer Transformer encoder acts as a discriminator. Through adversarial training, the discriminator compels the generator to learn not only the correct p-HLA-I pairing rules but also the complex sequence patterns inherent in the distribution of real HLA-I data. The detailed hyper-parameter settings of PSGM during the training process are presented in Supplementary Table S5. To conclude, for each generated pseudo-sequence, the mapping module employs a Hamming distance-based nearest-neighbor search algorithm to identify the most similar sequences within a reference database of 178 known pseudo sequences, outputting a ranked list of the closest HLA-I alleles.

### Pre-training Enhances Model Feature Representation

Prior to the primary training of the PHbinder model, we implemented a dedicated pre-training phase to enhance the model’s capacity for task-specific feature representation. This initial fine-tuning step, which integrates ESM2 with Low-Rank Adaptation (LoRA), allows the model to learn relevant latent patterns from the downstream data. This strategy not only provides a more effective parameter initialization for the main training phase but also significantly strengthens the feature extraction efficacy of the base ESM2 model for this specific immunological context.

The benefits of this two-stage training approach are evident in the model’s performance metrics (Figures 2a and 2b).As shown in Figure 2a, the pre-trained PHbinder model demonstrates a marked improvement across most metrics compared to a model trained from scratch. Specifically, it achieved increases of 0.43% in Accuracy, 0.32% in AUC, 0.62% in Precision, 1.66% in F1-score, and 0.01 in MCC, with only a marginal decrease in Recall. The training curves in Figure 2b further show that the two-stage strategy leads to faster convergence and earlier stabilization of performance on the validation set. Supplementary Tables S1 and S2 further respectively present in detail the performance of the validation set metrics and the decline in loss for each epoch with or without pre-training.

**Figure 2.**
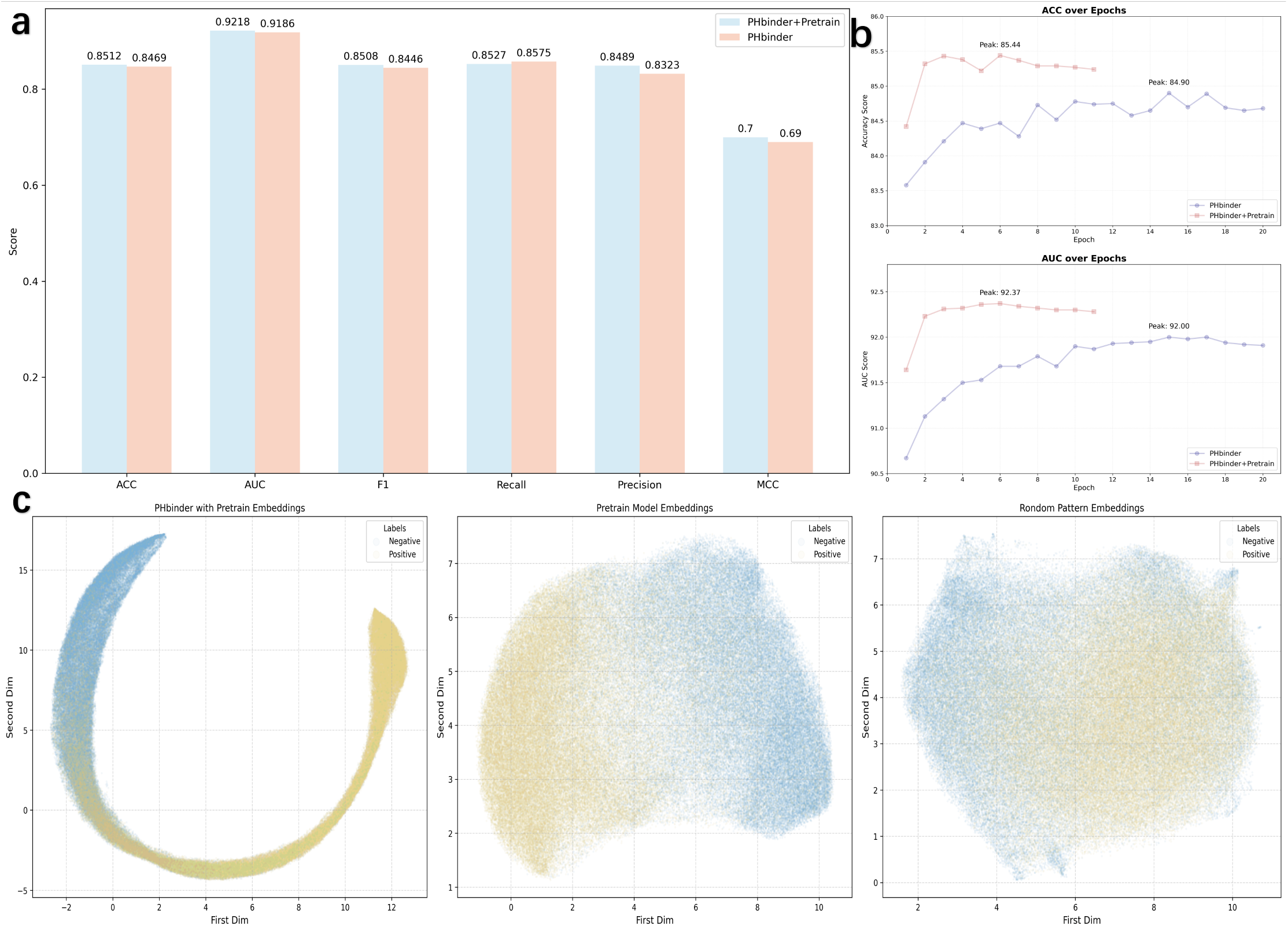
Performance comparison between PHbinder with pre-training (PHbinder+Pretrain) and without.a) Histogram comparing the final performance metrics of PHbinder with and without pre-training. b) Validation set AUC and MCC curves for PHbinder with and without pre-training across each training epoch, with the peak value and corresponding epoch annotated. c) UMAP distributions of positive (yellow) and negative (blue) samples at three stages: initial (raw ESM2), after LoRA pre-training, and after full training (PHbinder+Pretrain).

To visualize the impact of training on the feature space, we performed UMAP[29] dimensionality reduction on the sequence embeddings at different stages (Figure 2c). The analysis reveals a clear progression. In the raw, untrained ESM2 embeddings, positive and negative samples are indistinguishable and diffusely distributed. After the LoRA pre-training phase, the embeddings begin to coalesce into distinct groups, indicating emergent class separation. Following the full two-stage training of PHbinder, this separation becomes highly pronounced, with the samples forming well-defined and compact clusters. This visualization directly demonstrates that the discriminative power of the learned features was significantly enhanced throughout the training process.

### PHbinder Significantly Outperforms Baseline Sequence Classification Models

To validate the architectural design of PHbinder, we benchmarked its performance on the IEDB_HLA_I dataset against several standard sequence classification models, including TransHLA[16], TextCNN[30], DPCNN[31], RNN-ATT[32], and TextRCNN[33]. The task was framed as a binary classification problem: predicting whether a given peptide is an HLA-I binder.

The comparative results are presented in Table 1 and Figure 3b. PHbinder demonstrated superior performance, achieving the highest scores in four of the five key evaluation metrics: Accuracy (ACC) of 0.8512, F1-score of 0.8508, Matthews Correlation Coefficient (MCC) of 0.7024, and Area Under the ROC Curve (AUC) of 0.9218. Furthermore, PHbinder also led in Area Under the Precision-Recall Curve (AUPRC) with a score of 0.9185. In a direct comparison with the second-best model, TransHLA, PHbinder’s AUC and AUPRC were higher by 0.23% and 0.09%, respectively. While its recall was marginally lower than DPCNN’s, PHbinder’s overall performance profile was clearly superior. The consistent underperformance of simpler models, particularly TextCNN, suggests that capturing only local sequence patterns is insufficient for this task. We attribute PHbinder’s success to its hybrid architecture, which effectively integrates local features from the CNN branch with long-range dependency information from the Transformer branch. To assess the model’s generalization capabilities, we conducted further benchmark tests on three independent external datasets: IEDB_A, IEDB_B, and IEDB_C, which represent bindings to different HLA supertypes (Figure 3a). The ROC and PR curves in Figure 3b show that PHbinder consistently outperforms the baseline models across all three allele classes, confirming the robustness and general applicability of our approach.

**Table 1.** Performance Metrics of Various Models

| Model | ACC | F1 | Recall | MCC | AUC |
| --- | --- | --- | --- | --- | --- |
| PHbinder | 85.12 | 85.08 | 84.90 | 0.70 | 92.18 |
| TransHLA | 84.72 | 84.59 | 83.92 | 0.69 | 91.95 |
| TextCNN | 80.50 | 81.12 | 83.76 | 0.61 | 87.64 |
| DPCNN | 83.93 | 84.16 | 85.41 | 0.68 | 91.23 |
| RNN-ATTs | 82.66 | 82.65 | 82.64 | 0.65 | 90.01 |
| TextRCNN | 82.16 | 81.99 | 81.21 | 0.64 | 89.52 |

**Figure 3.**
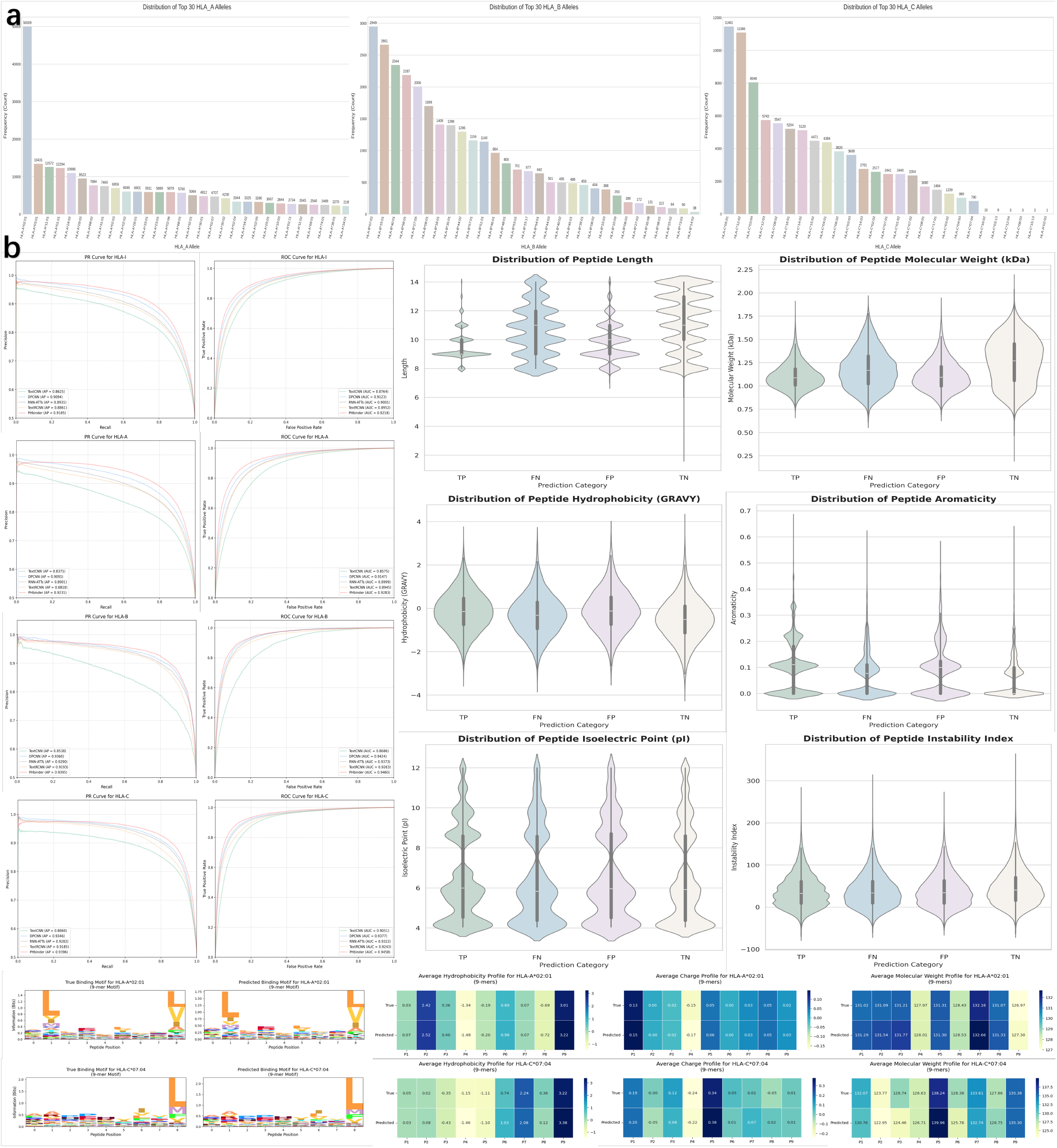
Performance comparison of PHbinder against baseline sequence classification models on various test sets.a) Distribution of the top 30 most frequent HLA-I alleles in the three external test sets: IEDB_A, IEDB_B, and IEDB_C. b) Precision-Recall (PR) and ROC curves for PHbinder and baseline models on the HLA_I, HLA_A, HLA_B, and HLA_C datasets, with corresponding AUC and AUPRC values. PHbinder comprehensively outperforms all baseline models. c) Violin plots illustrating six physicochemical properties of the peptides: length, molecular weight, hydrophobicity (GRAVY), aromaticity, isoelectric point (pI), and instability index. d) Sequence logos comparing true binding peptides with those predicted by PHbinder for HLA-A*02:11 and HLA-C*07:04 (for 9-mer peptides). e) Heatmaps showing the distribution of attention weights on physicochemical properties (hydrophobicity, charge, and molecular weight) at each peptide position for true vs. PHbinder-predicted binders of HLA-A*02:11 and HLA-C*07:04 (length 9).

Based on the frequency distribution in Figure 3a, we further analyzed the model’s performance on high-frequency and low-frequency alleles for peptides of varying lengths. Figures 3d and 3e show the prediction results for 9-mer peptides on a representative high-frequency allele (HLA-A*02:01) and a low-frequency allele (HLA-C*07:04).(Supplementary Figs.S1 shows the prediction results for peptides of other lengths.)For HLA-A*02:01, PHbinder identified the key P2 and P9 anchor positions and recognized the strong preference for specific hydrophobic amino acids at these sites (L at P2; V or L at P9). It also precisely reproduced the allele’s neutral charge characteristics and the exact spatial limitations of the binding groove. For HLA-C*07:04, the model accurately identified the main anchor position as P9 and predicted its preference for the hydrophobic amino acid Leucine (L), along with features of its strong electrostatic environment and per-position molecular weights. Supplementary Figs.S2 further shows the analysis on more high-frequency and low-frequency alleles. The results for these alleles across different peptide lengths, presented in Appendices 1 and 2, collectively show that the model learned the core anchor points and binding specificities for both high-frequency and low-frequency alleles.

Moreover, to investigate the features driving model predictions, we analyzed the physicochemical properties of peptides in each prediction category (TP, TN, FP, FN), visualized as violin plots in Figure 3c. This analysis revealed two key insights. First, the model is sensitive to physical properties like peptide length and molecular weight. Performance peaked for peptides of canonical length (9-10 amino acids), with misclassifications (particularly false negatives) increasing for non-standard lengths. Second, the model demonstrated remarkable robustness to chemical and structural properties. The distributions for hydrophobicity, aromaticity, isoelectric point, and instability were surprisingly consistent across all four prediction outcomes. This indicates that the model’s classifications are not based on simple biases toward specific chemical profiles but are instead driven by deeper, more complex sequence patterns learned during training.

### PHbinder Achieves State-of-the-Art Performance in Epitope Binding Prediction

To position PHbinder within the current landscape of immunoinformatics, we performed a comparative evaluation against six state-of-the-art p-HLA-I binding prediction tools: MHCflurry[24], TransPHLA[34], MixMHCpred[35], NetMHCpan-4.1b[36], Anthem[37], and MHCnuggets[38]. The assessment was conducted on the independent benchmark dataset, Immune_HLA_I. A fair comparison was ensured by converting the continuous outputs (e.g.,binding affinity or elution scores) from the benchmark tools into binary classification labels using their recommended thresholds, matching PHbinder’s output format.

The performance of all models on the Immune_HLA_I dataset is detailed in Table 2. The results show that PHbinder achieves state-of-the-art (SOTA) performance. With the exception of the Recall metric, where MHCnuggets achieved the highest value, PHbinder outperformed all other tools across three key metrics: Accuracy (0.8416), F1-score (0.8396), MCC (0.68). Compared to the second-best performing model in each respective category, PHbinder demonstrated improvements of 1.1% (ACC), 0.5% (F1), 2.9% (MCC). These results confirm that PHbinder’s comprehensive performance in predicting epitope binding to HLA class I molecules is superior to that of current leading methods.

**Table 2.** Performance Metrics of Different Methods

| Method | ACC | F1 | Recall | MCC | Precision | Specificity |
| --- | --- | --- | --- | --- | --- | --- |
| PHbinder | 84.16 | 83.96 | 82.90 | 0.68 | 85.05 | 85.43 |
| TransHLA | 83.09 | 82.56 | 80.06 | 0.66 | 85.22 | 86.11 |
| Mhcflurry | 82.96 | 83.24 | 84.57 | 0.66 | 81.95 | 81.38 |
| Mhc nuggets | 50.07 | 66.67 | 99.87 | 0.02 | 50.04 | 0.29 |
| TransPHLA | 62.68 | 71.75 | 94.78 | 0.33 | 57.72 | 30.58 |
| Anthem | 74.46 | 77.06 | 85.80 | 0.50 | 69.94 | 63.12 |
| NetMHCpan4.1b | 81.08 | 81.97 | 86.01 | 0.62 | 78.29 | 76.15 |
| MixMHCpred | 79.27 | 80.85 | 87.51 | 0.59 | 75.13 | 71.03 |

### Ablation Study of PHbinder

To dissect the architecture of PHbinder and quantify the contribution of its individual components, we conducted a series of six ablation studies. In these experiments, we systematically removed or replaced key modules to measure the impact on overall performance. The specific modifications were: 1) replacing Low-Rank Adaptation (LoRA) with traditional full-model fine-tuning; 2) replacing the pre-trained ESM2 encoder with a standard trainable embedding layer; 3) removing the CNN module; 4) removing the Transformer module; 5) replacing the Cross-Multi-Head Attention fusion mechanism with simple feature concatenation; and 6) replacing MimoLoss with a standard Cross-Entropy loss function.

The results, summarized in Table 3,Figure 4 and Supplementary Figs.S3, reveal the critical role of each component. The removal of the CNN module caused the most significant performance degradation, with the MCC and AUC dropping by 0.0321 and 1.84%, respectively.(Figure 4a) As visualized in the radar plot (Figure 4b), this configuration resulted in a comprehensive decline across all metrics, confirming that the local features captured by the CNN are indispensable. The second most impactful component was MimoLoss; its replacement led to a 0.0197 drop in MCC and a 1.43% drop in AUC, highlighting its effectiveness in shaping a better decision boundary. Replacing Cross-MultiHead Attention with simple concatenation also caused a substantial performance decrease (0.0168 in MCC, 0.85% in AUC), demonstrating its necessity for effectively fusing the heterogeneous local and global features.

**Table 3.**
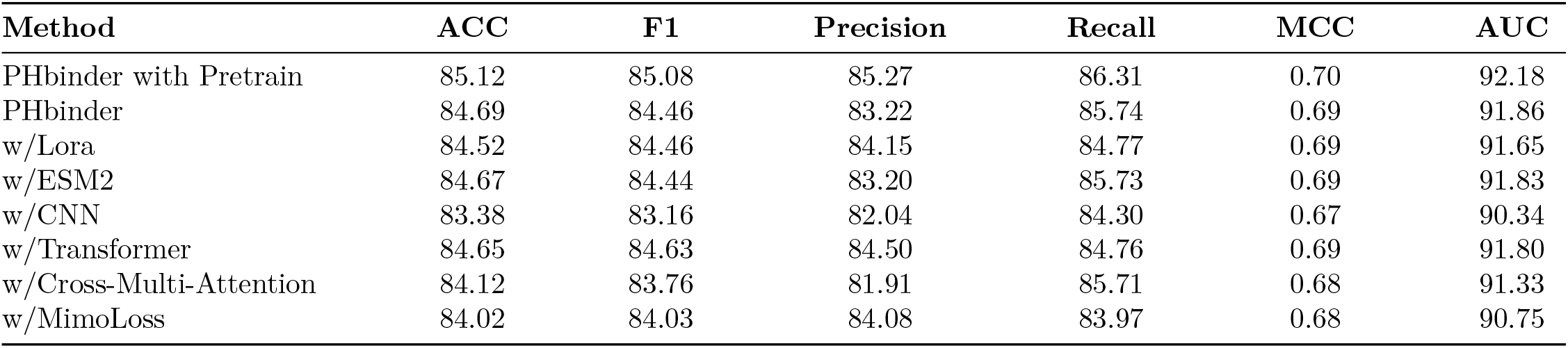
Ablation Study of PHbinder Components

| Method | ACC | F1 | Precision | Recall | MCC | AUC |
| --- | --- | --- | --- | --- | --- | --- |
| PHbinder with Pretrain | 85.12 | 85.08 | 85.27 | 86.31 | 0.70 | 92.18 |
| PHbinder | 84.69 | 84.46 | 83.22 | 85.74 | 0.69 | 91.86 |
| w/Lora | 84.52 | 84.46 | 84.15 | 84.77 | 0.69 | 91.65 |
| w/ESM2 | 84.67 | 84.44 | 83.20 | 85.73 | 0.69 | 91.83 |
| w/CNN | 83.38 | 83.16 | 82.04 | 84.30 | 0.67 | 90.34 |
| w/Transformer | 84.65 | 84.63 | 84.50 | 84.76 | 0.69 | 91.80 |
| w/Cross-Multi-Attention | 84.12 | 83.76 | 81.91 | 85.71 | 0.68 | 91.33 |
| w/MimoLoss | 84.02 | 84.03 | 84.08 | 83.97 | 0.68 | 90.75 |

**Table 4.**
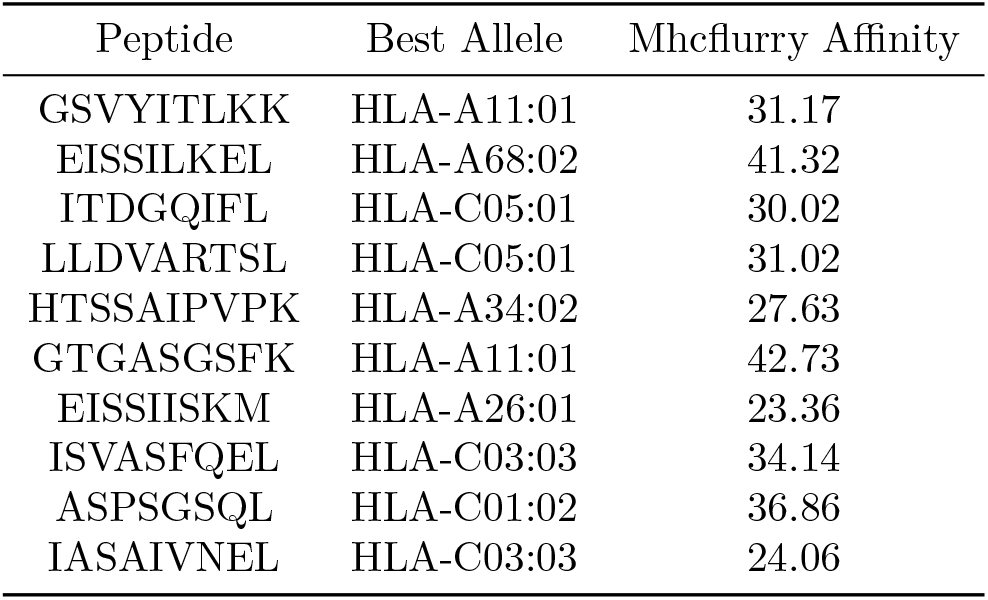
Potential p-HLA-I Binding

**Figure 4.**
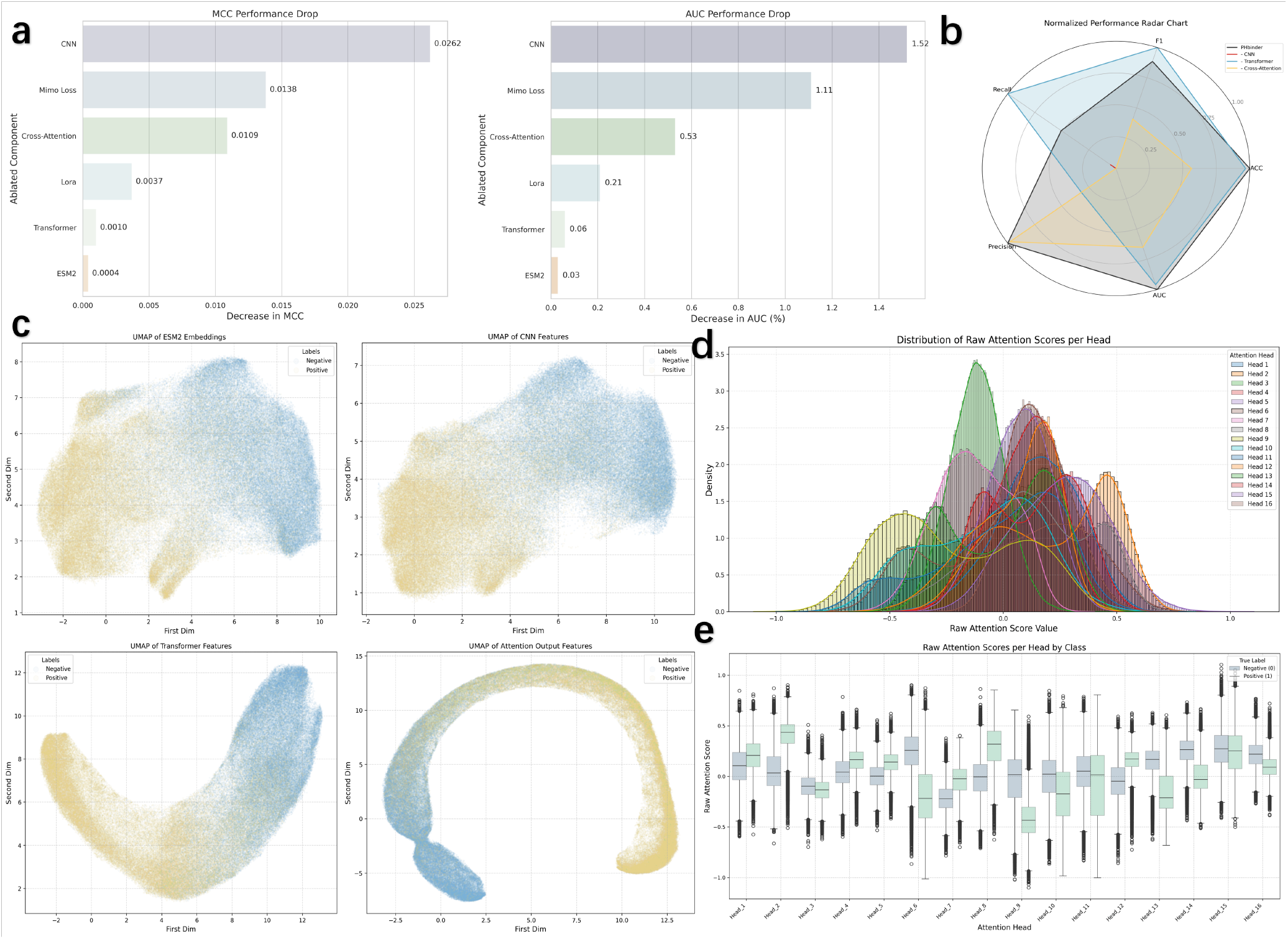
Ablation study of the PHbinder model. a) Histogram showing the performance degradation (in AUC and MCC) upon removing each of the six components: ESM2, LoRA, CNN, Transformer, Cross-Multi-Head Attention, and MimoLoss. b) Radar chart comparing the comprehensive performance of PHbinder after removing the CNN, Transformer, or Cross-Multi-Head Attention modules, respectively. c) UMAP visualization of features after extraction by ESM2, after processing by the CNN and Transformer modules, and after fusion by Cross-Multi-Attention. d) Distribution histograms of raw attention scores from the 16 heads of the Cross-Multi-Head Attention module. e) Box plots showing the attention score distributions for each of the 16 attention heads, separated by positive and negative class labels.

Furthermore, the comparison between LoRA and full fine-tuning yielded a noteworthy insight. Replacing LoRA with a full fine-tuning approach led to a clear, albeit smaller, decline in all metrics (e.g., a 0.0096 drop in MCC). This result indicates that for this task, the parameter-efficient LoRA technique is not only more computationally efficient but also serves as a more effective regularization strategy, leading to better generalization and superior overall performance.

To elucidate the model’s capacity to discriminate between phosphorylation sites, we visualized features from various training stages using Uniform Manifold Approximation and Projection (UMAP) (Figure 4c). The visualizations reveal a progressive refinement in class separability: initial features from ESM2 provide a baseline separation, which is markedly sharpened by the subsequent CNN and Transformer modules and ultimately optimized by the Cross-Multi-Head Attention fusion.

To dissect the internal strategy of the Cross-Multi-Head Attention module, we analyzed the raw attention scores from its 16 heads. Figure 4d displays the overall score distributions, while Figure 4e disaggregates these scores by their true labels. This analysis reveals a clear functional specialization among the heads developed during training. Notably, a subset of heads (6, 8, 9, and 15) exhibits potent class-discriminative capabilities, confirming that PHbinder successfully learns to identify and prioritize features most salient for the classification task.

### Performance Evaluation of PSGM on Known p-HLA-I Binding

To quantitatively evaluate the PSGM framework, we designed a rigorous protocol to assess its ability to generate biologically relevant HLA-I pseudo sequences and accurately map them to their correct binding partners. Using the IEDB_HLA_II dataset, we employed PSGM to autoregressively generate a pseudo sequence for each input epitope.

Supplementary Figs.S4 shows the frequency distribution of the dataset based on Allele classification. This generated sequence was then compared against a reference database of known HLA-I pseudo sequences using Hamming distance to produce a ranked list of candidate alleles.

We evaluated the generated sequences from three perspectives: sequence quality, distributional similarity, and novelty (Figure 5a). The model achieved a low Perplexity (PPL)[39] of 1.55 and a high average Sequence Recovery Rate (SRR) of 65.10%, indicating that the generated sequences are structurally coherent and closely resemble real sequences.

**Figure 5.**
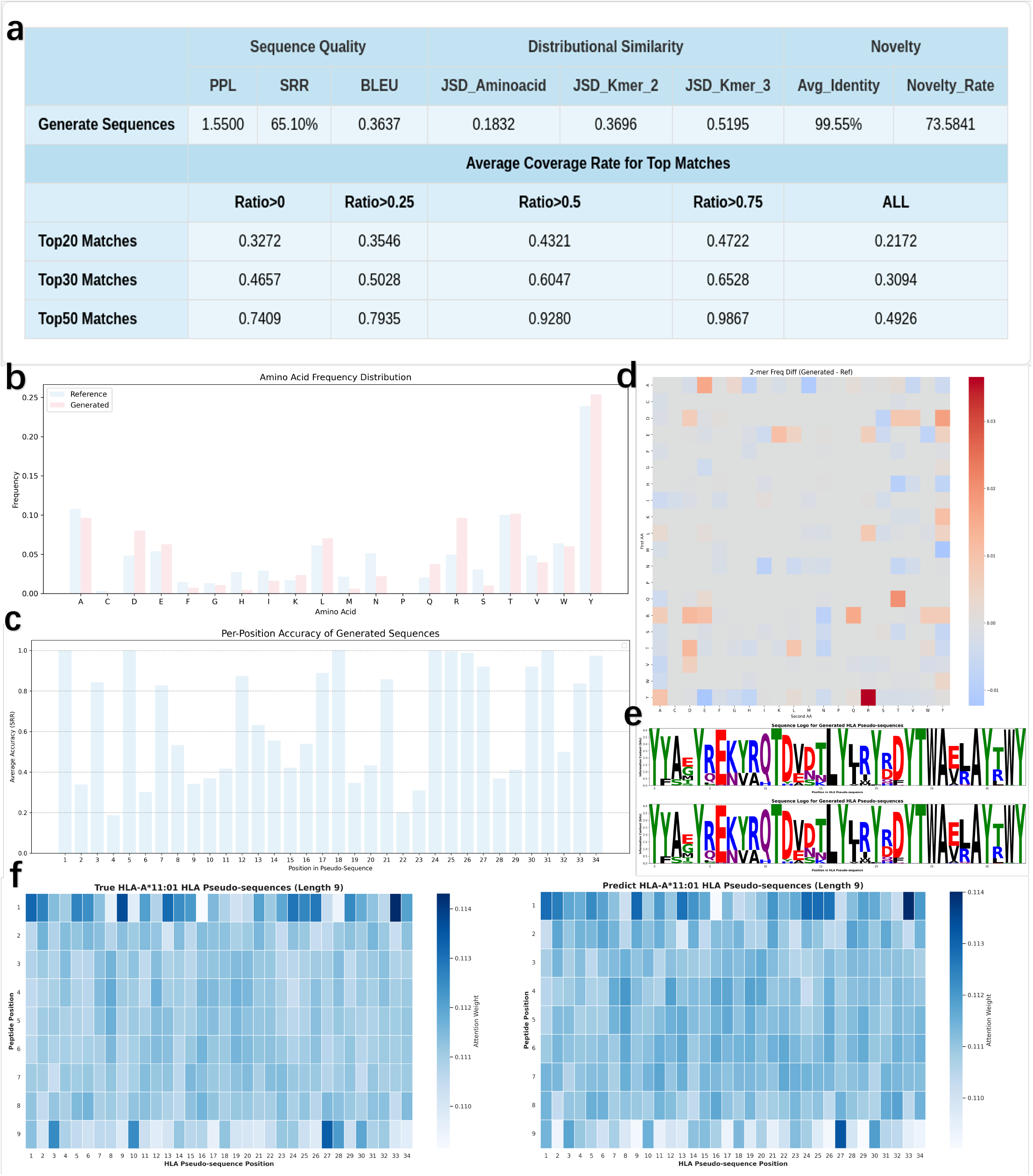
Comprehensive performance evaluation of the PSGM model. a) Performance evaluation of PSGM at both the generation and mapping levels. b) Amino acid frequency distribution plot. Compares the frequency of 20 amino acids between real sequences (blue) and generated sequences (red). c)Position-wise accuracy analysis. Displays the prediction accuracy for each position of the HLA-I pseudo sequence. d) Heatmap of 2-mer frequency differences. Illustrates the distributional difference between generated and real sequences for 2-mer amino acid fragments. The xand y-axes represent the first and second amino acid in the fragment, respectively. Gray indicates similarity. e) Sequence logos of authentic HLA-I pseudo-sequences versus those generated by PSGM. f) Attention distribution heatmaps for 9-mer peptide interactions, comparing authentic HLA-I pseudo-sequences with PSGM-generated ones for the HLA-A*11:01 allele.

This was corroborated at the local level by a BLEU[40] score of 0.3637. Figure 5c illustrates the accuracy rates at each amino acid position. Furthermore, analysis of Jensen-Shannon Divergence (JSD)[41] for amino acids (0.1832) and k-mers (0.3696 for 2-mers, 0.5195 for 3-mers) confirmed that the generated sequences faithfully replicate the statistical distributions found in real HLA-I data (Figures 5b and 5d). Critically, the model also demonstrated strong creative capacity. With a novelty rate of 99.55% (at a 90% sequence identity threshold) and an average sequence identity to the training set of only 73.58%, PSGM proves capable of generating a vast number of new, valid pseudo sequences while maintaining their core biological properties (Supplementary Figs.S5).

The primary metric for mapping performance was the Average Coverage Rate, defined as the proportion of experimentally validated HLA-I binders recovered within the model’s Top-k predictions, averaged across all test samples. As shown in Figure 5a, across all test entries, PSGM achieved average coverage rates of 21.72%, 30.94%, and 49.26% in its Top-20, Top-30, and Top-50 predictions, respectively. This demonstrates that, on average, the Top-50 list captures nearly half of all true binders for a given peptide. Performance was markedly higher for high-confidence predictions. For test cases where PSGM identified at least one correct binder (ratio > 0), the average coverage surged to 32.72% (Top-20), 46.57% (Top-30), and 74.09% (Top-50). In the most stringent scenario, for epitopes where the model recovered at least 75% of the true binders (ratio > 0.75), the average coverage in the Top-50 list reached an exceptional 98.67%, indicating a near-perfect recovery of known alleles. Additionally, we sought to illuminate the “black box” of PSGM by analyzing the weights learned by its attention layers, using HLA-A*11:01 as an example. As shown in Figure 5e, the sequence motifs of the pseudo-sequences generated by PSGM successfully capture the motifs of real HLA-I sequences at key anchor positions (including 0, 2, 12, 26, 30, 32, and 33).

Figure 5f and Supplementary Figs.S6 compare the attention distributions for peptides, contrasting the patterns from real HLA-peptide interactions in the training set with those from PSGM-generated interactions on an external dataset. This comparison, both at the aggregate and single-sample levels, shows that the model not only reproduces the overall attention patterns with high precision but also identifies the critical interaction sites between the peptide and the HLA-I pseudo-sequence. This indicates that PSGM genuinely focuses on the binding of key amino acids.

### PSGM Can Discover Potential p-HLA-I Binding

A significant limitation of evaluations based on known experimental data is their inherent incompleteness; many true binding pairs may be absent from databases and thus incorrectly treated as negatives. To overcome this bias, we performed an orthogonal validation using MHCflurry, a widely recognized independent prediction tool. This approach allowed us to assess the biological plausibility of PSGM’s novel predictions on the IEDB_Evaluate dataset, identifying potential p-HLA-I interactions that lack direct experimental support.

Our analysis revealed two key findings. First, PSGM’so prediction lists are highly enriched with viable binding candidates (Figures 6a and 6b). MHCflurry confirmed that 35.4% (Top-5), 35.0% (Top-10), 34.7% (Top-20), and 30.4% (Top-50) of the total predicted p-HLA-I pairs were likely binders. This high hit rate, averaging approximately one validated binder for every three predictions, demonstrates the quality of the candidate lists. Moreover, the remarkable stability of this enrichment as the list size increases underscores the robustness of PSGM’s predictive capability.

**Figure 6.**
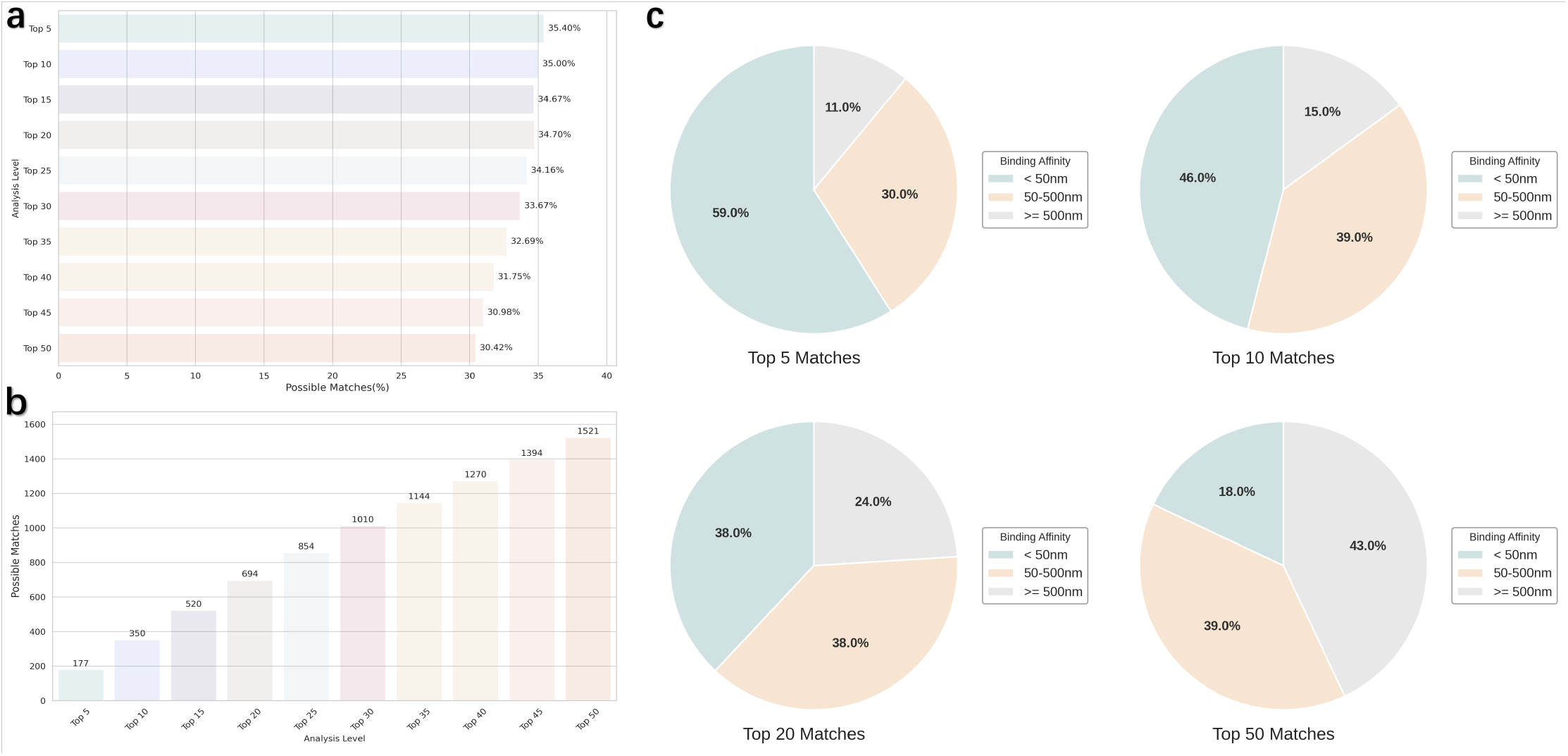
Validation of PSGM predictions using MHCflurry. a) Histogram showing the frequency distribution of MHCflurry-validated binding interactions among all alleles in the Top-50 prediction list. b) Horizontal bar plot showing the frequency of MHCflurry-validated binding for all alleles in the Top-50 list, depicting the overall predictive performance of PSGM. c) Pie charts of the best-matching allele validation status within the Top-5, Top-10, Top-20, and Top-50 prediction lists. Alleles are categorized as strong binders, weak binders, or non-binders based on binding affinity thresholds of 50 nM and 500 nM.

Second, and more importantly, PSGM demonstrated a powerful capacity for discovering novel binding pairs (Figure 6c). We focused on the single best-matching allele predicted by PSGM for each peptide within its Top-k lists. The validation rate for these top-ranked predictions was exceptionally high, with MHCflurry confirming 57% (Top-5), 68% (Top-10), 76% (Top-20), and 89% (Top-50) as high-affinity binders. Critically, a substantial fraction of these validated, high-confidence predictions were novel interactions not present in our training data. These newly discovered pairs constituted 9%, 22%, 26%, and 63% of the validated best matches in the Top-5, Top-20, Top-30, and Top-50 lists, respectively. Supplementary Table S3 provides a detailed presentation of these newly discovered binding pairs. Additionally, we used AlphaFold to verify whether these binding pairs can bind (Supplementary Figs.S7) This result confirms that PSGM not only reproduces known interactions but also effectively generalizes to identify novel p-HLA-I pairs that, while experimentally unverified, possess high predicted biological relevance.

## Discussion and conclusion

In this study, we developed a synergistic, two-stage framework to address critical gaps in p-HLA-I binding prediction. The first stage, PHbinder, functions as a high-throughput primary screen, leveraging a LoRA-finetuned ESM2 encoder and a hybrid CNN-Transformer architecture to accurately determine if a peptide is an immunogenic epitope. The second stage, PSGM, takes the epitopes identified by PHbinder and employs a generative model to predict a ranked list of their likely HLA-I allele binding partners. Our results demonstrate the success of this approach: PHbinder achieves state-of-the-art (SOTA) performance in epitope classification, and PSGM recovers, on average, 49.26% of known binding partners in its Top-50 predictions. This cascaded design transforms the conventional, labor-intensive “verification” workflow into an efficient “discovery” pipeline, establishing, for the first time, a direct “Peptide → Epitope Determination → HLA-I Alleles List” pathway.

A cornerstone of PHbinder’s success is the two-stage training strategy. By pre-training the LoRA-adapted ESM2 model on our task data before the final end-to-end training, we provided a superior parameter initialization. This initial phase allowed the model to develop a specific task feature space[42] attuned to the biochemical patterns of immunogenic peptides, significantly enhancing the performance of the final downstream classification task. Despite its strong performance, PHbinder has recognized limitations. The model was primarily designed for and validated on canonical linear peptides of 8-14 amino acids. Its performance on atypical antigens such as very long or short peptides, spliced peptides, or those with post-translational modifications (PTMs)[43] remains unverified. As these non-canonical epitopes are increasingly recognized as vital components of the antigenic landscape, future iterations of PHbinder will aim to broaden its applicability by incorporating dedicated feature channels to handle these complex cases.

Similarly, the performance of PSGM, while promising, exhibits variability. For some peptides, it can recover nearly all known binding alleles, whereas for others, its predictive power is limited. We attribute this instability primarily to two factors: 1) severe data imbalance in public immunopeptidomics datasets[44], which can bias the model towards common p-HLA-I pairs, and 2) our reliance on an implicit negative sampling strategy, where the absence of a recorded interaction is treated as a non-interaction. This assumption may introduce false negatives, constraining the model’s ability to generalize and discover novel binding patterns. We anticipate that as larger and more balanced datasets become available, the performance and stability of PSGM will improve significantly.

To validate the biological plausibility of PSGM’s novel predictions, we conducted an orthogonal in silico[45] evaluation using MHCflurry. This analysis confirmed that a high proportion of PSGM’s predicted novel p-HLA-I pairs had high binding affinities, indicating that our model has learned the underlying biochemical principles of molecular recognition, rather than merely memorizing statistical patterns in the training data.

In summary, the cascaded PHbinder-PSGM framework successfully reframes p-HLA-I binding analysis, moving it from a one-by-one “verification” task into a high-throughput “discovery” problem. This paradigm shift accelerates the screening of broad-spectrum vaccine candidates and deepens the analysis of immune escape mechanisms. Furthermore, our framework can serve as a powerful “upstream preprocessor” for traditional affinity predictors, focusing experimental efforts and enabling the discovery of novel, high-affinity p-HLA-I interactions.

## Experimental Section

### Datasets

#### IEDB_HLA_I

This is the core dataset for training and evaluating PHbinder. Sourced from the Immune Epitope Database (IEDB), it was curated by extracting all Class I HLA binding assays. The data was filtered for linear peptides (8-14 amino acids) with unambiguous positive or negative binding labels. After cleaning and deduplication, the dataset consists of 92,347 curated data points.

#### IEDB_A IEDB_B IEDB_C

These three datasets were created to test PHbinder’s performance on specific HLA supertypes. They consist of confirmed positive binding examples for HLA-A (449,320), HLA-B (55,942), and HLA-C (174,562) from IEDB, each balanced with an equal number of negative samples drawn from the NetMHCpan evaluation datasets.

#### Immune_HLA_I

This independent external validation set was constructed to benchmark PHbinder against other state-of-the-art (SOTA) tools. It integrates 21,387 experimentally verified p-HLA-I pairs curated from multiple public databases, including CEDAR[46], VDJdb[47], ImmunoCODE, and dbPepNeo2.0[48].

#### IEDB_HLA_II

This dataset was specifically curated to train the PSGM generative model. It was sourced from IEDB and contains only experimentally confirmed positive binding interactions. This “positive-only” design provides PSGM with highquality exemplars of p-HLA-I binding rules. The final dataset contains 341,713 p-HLA-I pairs where the peptide is 8-14 amino acids in length.

#### Pseudo_HLA

This reference database was constructed for the mapping module of the PSGM framework. The creation process was as follows: 1) All HLA-I alleles from IEDB_HLA_II were compiled. 2) Their full-length amino acid sequences were retrieved from the IMGT/HLA database. 3) The 34 key amino acid residues involved in peptide binding were extracted to form a “pseudo sequence” for each allele, following the NetMHCpan definition. 4) Since multiple alleles can share the same pseudo sequence, a single canonical allele was chosen as the representative for each unique pseudo sequence to resolve this many-to-one mapping. 5) We constructed a mapping database containing 178 unique HLA-I pseudo sequences, representing over 10,000 known HLA-I alleles.

#### IEDB_Evaluate

This independent test set was designed for the orthogonal validation of PSGM’s novel predictions. It contains 100 unique peptides, sourced from public databases like CEDAR, that were absent from all training sets. This dataset is used to generate candidate HLA-I allele lists with PSGM, which are then evaluated for biological plausibility by the independent MHCflurry tool.

### LoRA(Low-Rank Adaptation)

Large pre-trained language models (such as ESM-2) incur substantial computational and storage costs when undergoing full fine-tuning for down-stream tasks. To achieve parameter-efficient tuning, we employed the Low-Rank Adaptation (LoRA) strategy [19]. The core idea of LoRA is that the change in the model’s parameter matrix when adapting to a new task is low-rank. Therefore, instead of directly updating the pre-trained weight matrix **W**_0_ ∈ ℝ^*d×k*^, we introduce a bypass structure composed of two low-rank matrices to represent its update Δ**W**.

Specifically, we decompose the weight update Δ**W** into the product of two smaller matrices:

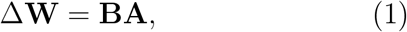

where **B** ∈ ℝ^*d×r*^ and **A** ∈ ℝ^*r×k*^ . Here, *r* is a rank much smaller than *d* and *k*, i.e., *r ≪* min(*d, k*). During training, the pre-trained weights **W**_0_ remain frozen and only the matrices **A** and **B** are trainable. For an input vector **x**, the modified forward propagation process can be expressed as:

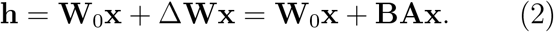

Matrix **A** is typically initialized using a random Gaussian distribution, while matrix **B** is initialized as a zero matrix. This ensures that at the beginning of training, Δ**W** = **BA** is zero, keeping the model in its original pre-trained state. Additionally, we apply a scaling factor 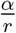 to the output of **BAx**, where *α* is a constant hyperparameter. This approach significantly reduces the number of parameters that need to be trained (from *d × k* to *r ×* (*d*+*k*)) while demonstrating comparable or even superior performance to full fine-tuning in downstream tasks.

### Transformer Encoder

To capture long-range dependencies between amino acid residues in epitope sequences, we employ the standard transformer encoder architecture [17]. This encoder consists of *N* = 6 identical layers stacked on top of each other. Each encoder layer comprises two core submodules: Multi-Head Self-Attention and Position-wise Feed-Forward Network.

For an input sequence embedding 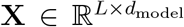, where *L* is the sequence length and *d*_model_ is the embedding dimension (640 in our model), the self-attention mechanism first linearly projects it into three distinct spaces: Queries (**Q**), Keys (**K**), and Values (**V**):

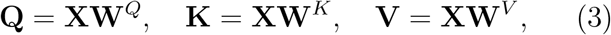

where 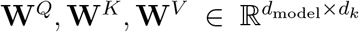 are learnable projection matrices. The attention scores are computed by taking the dot product of the Queries with all Keys, followed by normalization with a scaling factor 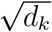 to prevent vanishing gradients. Finally, the softmax function is applied to obtain the attention weights, which are used to weight sum the Value vectors:

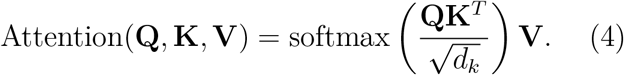

In the multi-head attention mechanism, this process is executed in parallel *h* times (16 in our model), each with distinct projection matrices. The outputs from each head are concatenated and subjected to another linear projection to produce the final output. This mechanism allows the model to jointly attend to information from different positions across various representational subspaces.

### Convolutional Neural Network Module

Unlike the Transformer’s ability to capture global dependencies, Convolutional Neural Networks (CNNs) excel at extracting local patterns and position-invariant features from sequences, which is crucial for identifying conserved immunogenic motifs. Our CNN branch consists of a series of one-dimensional convolutional layers and residual connections.

Given the sequence embeddings 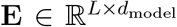 obtained from ESM-2, we first transpose the channel dimension to obtain 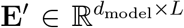. The first convolutional layer `region_cnn1` uses a kernel size of 3 to map the *d*_model_-dimensional input channels to *c*_out_ = 256 output channels, which can be represented as:

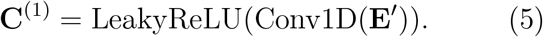

Subsequently, the feature map **C**^(1)^ enters a deep residual CNN block `cnn_block2`. This block first applies max-pooling to reduce the sequence dimension and enhance the feature’s translational invariance. Then, it extracts features through two consecutive 1D convolutional layers (`cnn1`) and adds them residually to the pooled input. This operation can be expressed as:

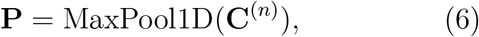

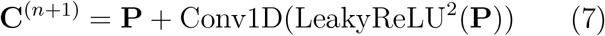

By stacking multiple such residual blocks, the CNN branch can effectively learn hierarchical features ranging from local to semi-global.

### Cross-Multi-Attention Mechanism

To effectively integrate the global contextual information captured by the Transformer branch and the local motif features extracted by the CNN branch, we designed a cross-attention module. This mechanism allows for asymmetric interaction between feature representations from two different sources.

In this module, we take the final output feature vector 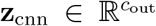 from the CNN branch as the query (Query), and the global representation vector 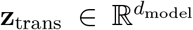 from the Transformer branch as both the key (Key) and value (Value). Initially, these three vectors are projected into the subspace of the multi-head attention:

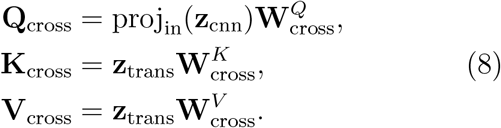

Here, proj_in_ is an initial convolutional layer used to align the dimensions of **z**_cnn_. Note that the query originates from local features, while both the key and value stem from global features. The computation of attention weights is similar to self-attention but reflects the correlation between local and global features:

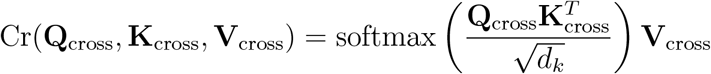

By doing so, the model learns how to dynamically extract and integrate the most relevant information from the global context based on a given local motif (query), thereby generating a more informative and discriminative fused feature representation for the final classification task.

### MimoLoss

In our model, to optimize the training process and enhance the model’s generalization capability and robustness, we designed and adopted a composite loss function named MimoLoss (Mutual Information Maximization Loss). The design of this loss function draws on concepts from information theory, combining standard cross-entropy loss with entropy regularization and conditional entropy constraints, aiming to guide the model to learn more discriminative feature representations.

The core of MimoLoss consists of three parts. First, we calculate a modified cross-entropy loss 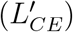, which serves as the basis for measuring the difference between the model’s predicted distribution and the true label distribution. Unlike standard cross-entropy, we introduce a small offset term, in the form of (loss - 0.04).abs() + 0.04, which helps stabilize gradients during the initial training phase and does not penalize minor prediction errors. Secondly, to enhance the model’s robustness, we introduce an entropy regularization term. This term calculates the entropy 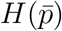 of the average predicted probabilities across the entire batch. By maximizing this term, we encourage the model to produce diverse predictions across different samples, preventing the model from falling into trivial solutions (e.g., predicting the same class for all inputs), thereby enhancing the model’s exploratory capabilities.

Lastly, we incorporate a conditional entropy constraint term *H*(*Y* | *X*). This term calculates the entropy of each sample’s predicted probability distribution and takes the average across the entire batch. By minimizing the conditional entropy, we encourage the model to make high-confidence (i.e., low uncertainty) predictions for each individual sample. This prompts the model to learn deterministic features that clearly separate different classes. Combining the above three parts, the final mathematical expression of MimoLoss is as follows:

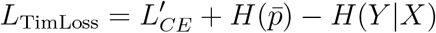

where: 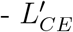 is the modified cross-entropy loss. 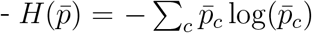 is the entropy of the batch-averaged predictions, where 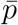 is the mean of the predicted probabilities for all samples in the batch. 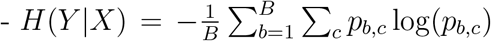 is the conditional entropy of predictions, with *B* being the batch size.

In this way, MimoLoss not only ensures that the model fits the training data but also regularizes the model by maximizing prediction diversity and minimizing predictive uncertainty, thereby effectively enhancing the model’s performance and generalization capability in complex classification tasks such as protein antigen recognition.

### Generative Adversarial Network

To address the complex task of generating corresponding HLA pseudo sequences for a given immunogenic peptide under the condition, we conducted “Generative Adversarial Network” (GAN) framework. The GAN framework consists of two core, mutually antagonistic sub-networks: a conditional sequence generator (Generator, *G*) and a sequence discriminator (Discriminator, *D*).

The generator *G* aims to learn the mapping from a conditional peptide sequence **P** = (*p*_1_, *p*_2_, …, *p*_*m*_) to the target HLA pseudo sequence **H** = (*h*_1_, *h*_2_, …, *h*_*T*_). It is constructed as a Transformer-based encoderdecoder model capable of capturing contextual information from the input peptide and autoregressively generating the output sequence. Its core function is to estimate the conditional probability distribution of the next amino acid *h*_*t*_ given the conditional **P** and the previously generated prefix **H**_*<t*_:

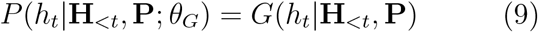

where *θ*_*G*_ represents the learnable parameters of the generator *G*. To obtain high-quality conditional representations, the generator’s encoding part utilizes a pre-trained ESM-2 protein language model to extract deep semantic features from the peptide sequence **P**, forming a conditional memory state **M**. The decoder then combines this memory state **M** with the autoregressive generation of the HLA sequence through a cross-attention mechanism, ensuring that the generation process strictly follows the guidance of the input condition. The discriminator *D* serves to evaluate the authenticity of a given HLA pseudo sequence **H**, i.e., determining whether it originates from the real dataset *p*_data_ or is synthesized by the generator *G*. It is designed as a Transformer-based sequence classifier capable of mapping the input sequence **H** to a scalar probability value *D*(**H**; *θ*_*D*_) ∈ [0, 1], where *θ*_*D*_ are its learnable parameters. This probability value indicates the likelihood that **H** is a genuine sequence.

The training process of the GAN framework is a minimax game driven by a unified objective function comprising two parts. The generator *G*’s objective function *L*_*G*_ aims to minimize a composite loss consisting of traditional supervised learning loss and adversarial loss:

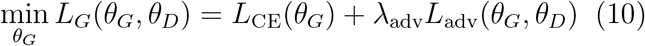

where *L*_CE_ is the standard cross-entropy loss (or negative log-likelihood) used to maximize the probability of generating the true target sequence **H** given the condition **P**. It ensures the accuracy of the generated content:

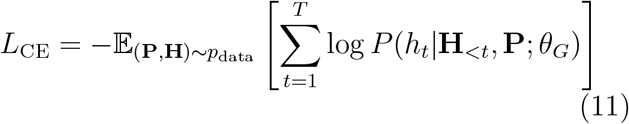

*L*_adv_ is the adversarial loss that encourages the generator *G* to produce sequences that can “deceive” the discriminator *D*, i.e., maximizing the probability that *D* misclassifies the generated samples as genuine. This part of the loss is used to adversarially refine the generation, making it statistically closer to real data:

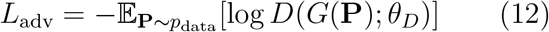

Meanwhile, the discriminator *D*’s training objective is to maximize its classification accuracy, i.e., minimizing the following loss function *L*_*D*_:

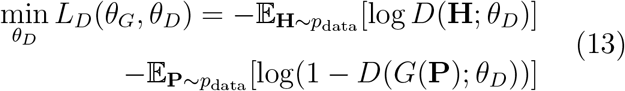

By alternatingly optimizing these two interrelated objective functions, the GAN model not only learns the precise mapping rules from peptide to HLA pseudo sequence but also, through the adversarial training process, learns the intrinsic syntax and high-dimensional distributional features of genuine HLA pseudo sequences, thus being able to generate sequences that are both accurate and highly realistic.

## References

[1] Charles Janeway, Paul Travers, Mark Walport, Mark J Shlomchik, et al. Immunobiology: the immune system in health and disease, volume 2. Garland Pub. New York, NY, USA, 2001.

[2] ARM Townsend, J Rothbard, FM Gotch, G Bahadur, D Wraith, and AJ McMichael. The epitopes of influenza nucleoprotein recognized by cytotoxic t lymphocytes can be defined with short synthetic peptides. Cell, 44(6):959–968, 1986.

[3] Mark M Davis and Pamela J Bjorkman. T-cell antigen receptor genes and t-cell recognition. Nature, 334(6181):395–402, 1988.

[4] K Christopher Garcia, Luc Teyton, and Ian A Wilson. Structural basis of t cell recognition. Annual review of immunology, 17(1):369–397, 1999.

[5] Gavin P Dunn, Allen T Bruce, Hiroaki Ikeda, Lloyd J Old, and Robert D Schreiber. Cancer immunoediting: from immunosurveillance to tumor escape. Nature immunology, 3(11):991– 998, 2002.

[6] Rolf M Zinkernagel and Peter C Doherty. Restriction of in vitro t cell-mediated cytotoxicity in lymphocytic choriomeningitis within a syngeneic or semiallogeneic system. Nature, 248(5450):701–702, 1974.

[7] Benoît J Van den Eynde and T Boon. Tumor antigens recognized by t lymphocytes. International Journal of Clinical and Laboratory Research, 27:81–86, 1997.

[8] Ugur Sahin, Evelyna Derhovanessian, Matthias Miller, Björn-Philipp Kloke, Petra Simon, Martin Löwer, Valesca Bukur, Arbel D Tadmor, Ulrich Luxemburger, Barbara Schrörs, et al. Personalized rna mutanome vaccines mobilize poly-specific therapeutic immunity against cancer. Nature, 547(7662):222– 226, 2017.

[9] Bjoern Peters, Morten Nielsen, and Alessandro Sette. T cell epitope predictions. Annual Review of Immunology, 38(1):123–145, 2020.

[10] Gary D Stormo. Dna binding sites: representation and discovery. Bioinformatics, 16(1):16–23, 2000.

[11] Kenneth C Parker, Maria A Bednarek, and John E Coligan. Scheme for ranking potential hla-a2 binding peptides based on independent binding of individual peptide sidechains. Journal of immunology (Baltimore, Md.: 1950), 152(1):163–175, 1994.

[12] H-G Rammensee, Jutta Bachmann, Niels Philipp Nikolaus Emmerich, Oskar Alexander Bachor, and Ssyfpeithi Stevanovic. Syfpeithi: database for mhc ligands and peptide motifs. Immunogenetics, 50:213–219, 1999.

[13] Morten Nielsen, Claus Lundegaard, Thomas Blicher, Kasper Lamberth, Mikkel Harndahl, Sune Justesen, Gustav Røder, Bjoern Peters, Alessandro Sette, Ole Lund, et al. Netmhcpan, a method for quantitative predictions of peptide binding to any hla-a and-b locus protein of known sequence. PloS one, 2(8):e796, 2007.

[14] Timothy J O’Donnell, Alex Rubinsteyn, Maria Bonsack, Angelika B Riemer, Uri Laserson, and Jeff Hammerbacher. Mhcflurry: opensource class i mhc binding affinity prediction. Cell systems, 7(1):129–132, 2018.

[15] Donald F Hunt, Robert A Henderson, Jeffrey Shabanowitz, Kazuyasu Sakaguchi, Hanspeter Michel, Noelle Sevilir, Andrea L Cox, Ettore Appella, and Victor H Engelhard. Characterization of peptides bound to the class i mhc molecule hla-a2. 1 by mass spectrometry. Science, 255(5049):1261–1263, 1992.

[16] Tianchi Lu, Xueying Wang, Wan Nie, Miaozhe Huo, and Shuaicheng Li. Transhla: a hybrid transformer model for hla-presented epitope detection. GigaScience, 14:giaf008, 2025.

[17] Vaswani Ashish. Attention is all you need. Advances in neural information processing systems, 30:I, 2017.

[18] Hezheng Lin, Xing Cheng, Xiangyu Wu, and Dong Shen. Cat: Cross attention in vision transformer. In 2022 IEEE international conference on multimedia and expo (ICME), pages 1–6. IEEE, 2022.

[19] Edward J Hu, Yelong Shen, Phillip Wallis, Zeyuan Allen-Zhu, Yuanzhi Li, Shean Wang, Lu Wang, Weizhu Chen, et al. Lora: Low-rank adaptation of large language models. ICLR, 1(2):3, 2022.

[20] Zeming Lin, Halil Akin, Roshan Rao, Brian Hie, Zhongkai Zhu, Wenting Lu, Nikita Smetanin, Robert Verkuil, Ori Kabeli, Yaniv Shmueli, et al. Evolutionary-scale prediction of atomic-level protein structure with a language model. Science, 379(6637):1123–1130, 2023.

[21] Ian J Goodfellow, Jean Pouget-Abadie, Mehdi Mirza, Bing Xu, David Warde-Farley, Sherjil Ozair, Aaron Courville, and Yoshua Bengio. Generative adversarial nets. Advances in neural information processing systems, 27, 2014.

[22] Richard W Hamming. Error detecting and error correcting codes. The Bell system technical journal, 29(2):147–160, 1950.

[23] Jon Louis Bentley. Multidimensional binary search trees used for associative searching. Communications of the ACM, 18(9):509–517, 1975.

[24] Timothy J O’Donnell, Alex Rubinsteyn, and Uri Laserson. Mhcflurry 2.0: improved pan-allele prediction of mhc class i-presented peptides by incorporating antigen processing. Cell systems, 11(1):42–48, 2020.

[25] Bent Fuglede and Flemming Topsoe. Jensenshannon divergence and hilbert space embedding. In International symposium on Information theory, 2004. ISIT 2004. Proceedings., page 31. IEEE, 2004.

[26] Carl H June and Michel Sadelain. Chimeric antigen receptor therapy. New England Journal of Medicine, 379(1):64–73, 2018.

[27] Randi Vita, Swapnil Mahajan, James A Overton, Sandeep Kumar Dhanda, Sheridan Martini, Jason R Cantrell, Daniel K Wheeler, Alessandro Sette, and Bjoern Peters. The immune epitope database (iedb): 2018 update. Nucleic acids research, 47(D1):D339–D343, 2019.

[28] Sumudu P Leelananda and Steffen Lindert. Computational methods in drug discovery. Beilstein journal of organic chemistry, 12(1):2694–2718, 2016.

[29] Leland McInnes, John Healy, and James Melville. Umap: Uniform manifold approximation and projection for dimension reduction. arXiv preprint 1802.03426, 2018.

[30] Yahui Chen. Convolutional neural network for sentence classification. Master’s thesis, University of Waterloo, 2015.

[31] Rie Johnson and Tong Zhang. Deep pyramid convolutional neural networks for text categorization. In Proceedings of the 55th Annual Meeting of the Association for Computational Linguistics (Volume 1: Long Papers), pages 562–570, 2017.

[32] Zichao Yang, Diyi Yang, Chris Dyer, Xiaodong He, Alex Smola, and Eduard Hovy. Hierarchical attention networks for document classification. In Proceedings of the 2016 conference of the North American chapter of the association for computational linguistics: human language technologies, pages 1480–1489, 2016.

[33] Siwei Lai, Liheng Xu, Kang Liu, and Jun Zhao. Recurrent convolutional neural networks for text classification. In Proceedings of the AAAI conference on artificial intelligence, volume 29, 2015.

[34] Yanyi Chu, Yan Zhang, Qiankun Wang, Lingfeng Zhang, Xuhong Wang, Yanjing Wang, Dennis Russell Salahub, Qin Xu, Jianmin Wang, Xue Jiang, et al. A transformer-based model to predict peptide–hla class i binding and optimize mutated peptides for vaccine design. Nature Machine Intelligence, 4(3):300–311, 2022.

[35] Daniel M Tadros, Julien Racle, and David Gfeller. Predicting mhc-i ligands across alleles and species: how far can we go? Genome medicine, 17(1):25, 2025.

[36] Birkir Reynisson, Bruno Alvarez, Sinu Paul, Bjoern Peters, and Morten Nielsen. Netmhcpan-4.1 and netmhciipan-4.0: improved predictions of mhc antigen presentation by concurrent motif deconvolution and integration of ms mhc eluted ligand data. Nucleic acids research, 48(W1):W449–W454, 2020.

[37] Shutao Mei, Fuyi Li, Dongxu Xiang, Rochelle Ayala, Pouya Faridi, Geoffrey I Webb, Patricia T Illing, Jamie Rossjohn, Tatsuya Akutsu, Nathan P Croft, et al. Anthem: a user customised tool for fast and accurate prediction of binding between peptides and hla class i molecules. Briefings in Bioinformatics, 22(5):bbaa415, 2021.

[38] Xiaoshan M Shao, Rohit Bhattacharya, Justin Huang, IK Ashok Sivakumar, Collin Tokheim, Lily Zheng, Dylan Hirsch, Benjamin Kaminow, Ashton Omdahl, Maria Bonsack, et al. High-throughput prediction of mhc class i and ii neoantigens with mhcnuggets. Cancer immunology research, 8(3):396–408, 2020.

[39] Fred Jelinek, Robert L Mercer, Lalit R Bahl, and James K Baker. Perplexity—a measure of the difficulty of speech recognition tasks. The Journal of the Acoustical Society of America, 62(S1):S63–S63, 1977.

[40] Kishore Papineni, Salim Roukos, and T Ward. Zhu w.-j. 2002. bleu: a method for automatic evaluation of machine translation. In Proceedings of the 40th annual meeting of the association for computational linguistics, volume 10, 1997.

[41] María Luisa Menéndez, Julio Angel Pardo, Leandro Pardo, and María del C Pardo. The jensen-shannon divergence. Journal of the Franklin Institute, 334(2):307–318, 1997.

[42] Jacques Neefjes, Marlieke LM Jongsma, Petra Paul, and Oddmund Bakke. Towards a systems understanding of mhc class i and mhc class ii antigen presentation. Nature reviews immunology, 11(12):823–836, 2011.

[43] Yoshua Bengio, Aaron Courville, and Pascal Vincent. Representation learning: A review and new perspectives. IEEE transactions on pattern analysis and machine intelligence, 35(8):1798–1828, 2013.

[44] James Robinson, Jason A Halliwell, James D Hayhurst, Paul Flicek, Peter Parham, and Steven GE Marsh. The ipd and imgt/hla database: allele variant databases. Nucleic acids research, 43(D1):D423–D431, 2015.

[45] S Ekins, J Mestres, and B Testa. In silico pharmacology for drug discovery: applications to targets and beyond. British journal of pharmacology, 152(1):21–37, 2007.

[46] Raghupathy Sivakumar, Prasun Sinha, and Vaduvur Bharghavan. Cedar: a core-extraction distributed ad hoc routing algorithm. IEEE Journal on Selected Areas in communications, 17(8):1454–1465, 2002.

[47] Dmitry V Bagaev, Renske MA Vroomans, Jerome Samir, Ulrik Stervbo, Cristina Rius, Garry Dolton, Alexander Greenshields-Watson, Meriem Attaf, Evgeny S Egorov, Ivan V Zvyagin, et al. Vdjdb in 2019: database extension, new analysis infrastructure and a t-cell receptor motif compendium. Nucleic acids research, 48(D1):D1057–D1062, 2020.

[48] Manman Lu, Linfeng Xu, Xingxing Jian, Xiaoxiu Tan, Jingjing Zhao, Zhenhao Liu, Yu Zhang, Chunyu Liu, Lanming Chen, Yong Lin, et al. dbpepneo2. 0: A database for human tumor neoantigen peptides from mass spectrometry and tcr recognition. Frontiers in Immunology, 13:855976, 2022.

