## Supplementary Materials for "PHbinder and PSGM: A Cascaded Framework for Epitope Prediction and HLA-I Allele Identification"

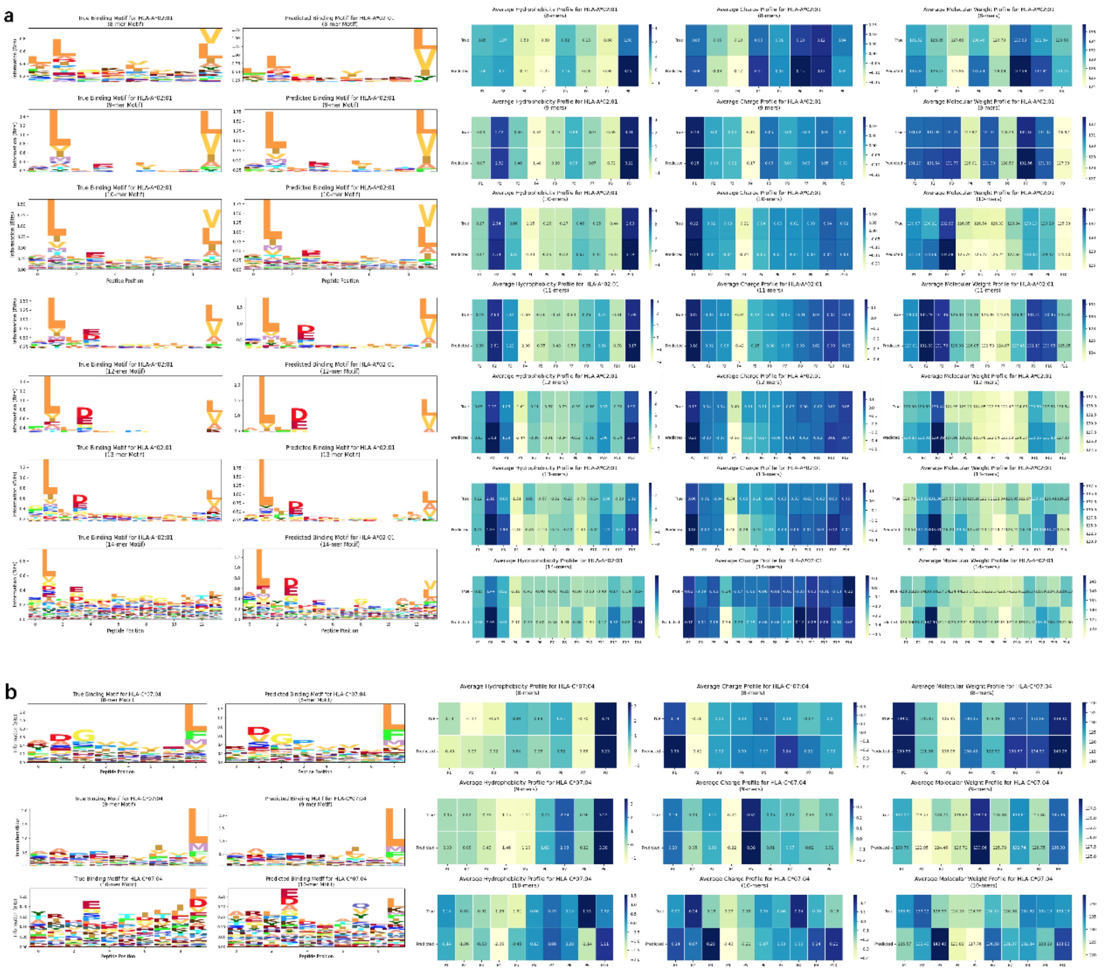


**Figure S1.** The prediction performance of HLA-A*02:01 and HLA-C*07:04 on peptide segments of various lengths is respectively presented (a detailed analysis of the 9-mer peptide segment situation has been provided in the main text). The reason why only some length cases are shown in Figure b is that we set a minimum threshold of 10 peptide segments for each category. The left side is the sequence logo plot made on peptide segments of various lengths, reflecting the comparison of the real sequence motifs and the predicted sequence motifs at each position. The right side is the comparison of physicochemical properties at each position, from left to right being hydrophobicity, charge distribution, and molecular weight.

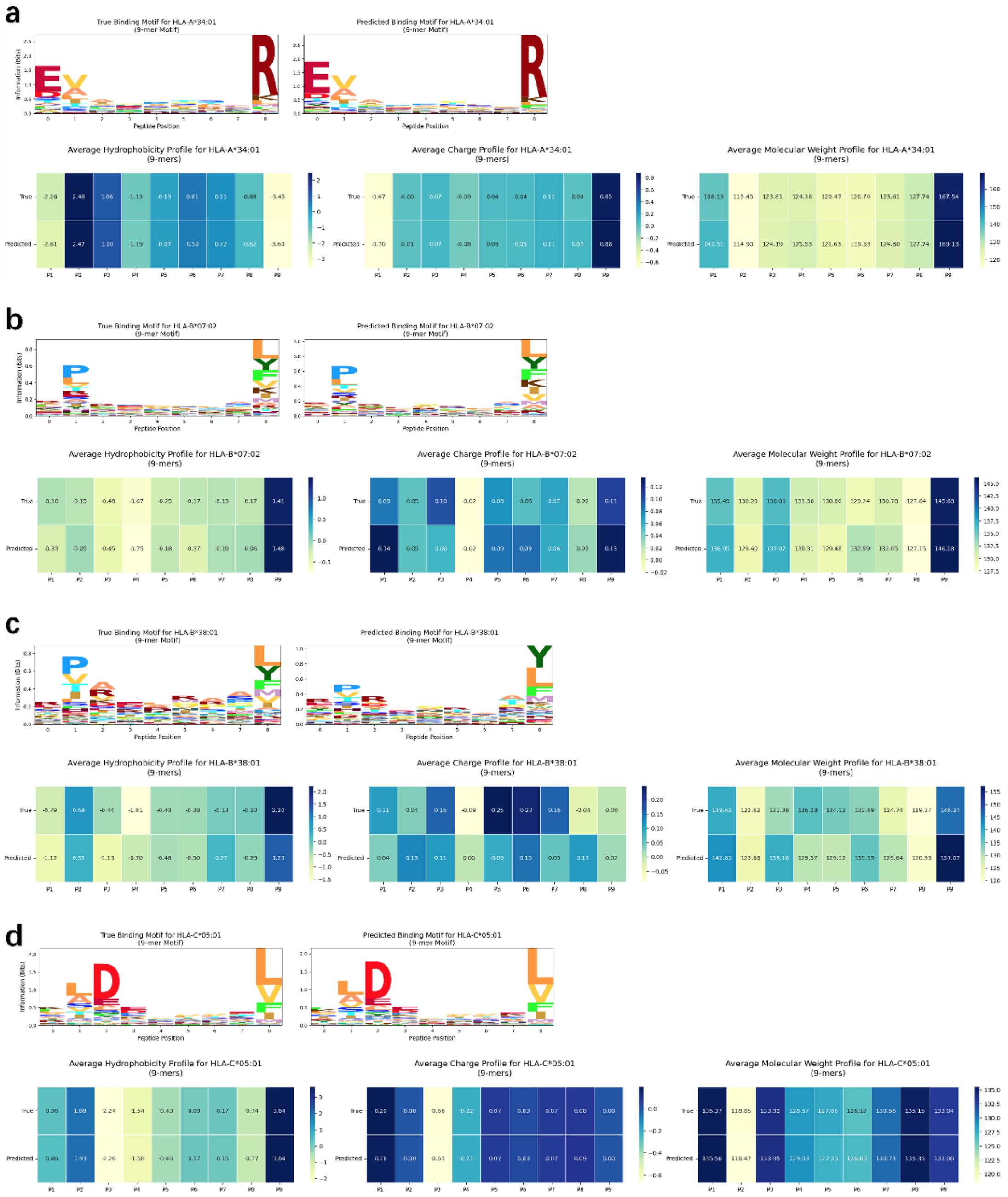


**Figure S2.** The prediction results of specific alleles on 9-mer peptides under various high-frequency and low-frequency cases were respectively presented. We selected the alleles with the highest and lowest frequency distribution (with at least 100 peptide segment source data for the lowest) in each HLA-A, HLA-B, and HLA-C dataset for detailed analysis to comprehensively reflect the prediction performance of PHbinder under different alleles. Each subfigure consists of two parts: the sequence logo plot and the physicochemical property attention distribution plot, which reflect the sequence motifs at each position and the analysis and verification at the physicochemical level.

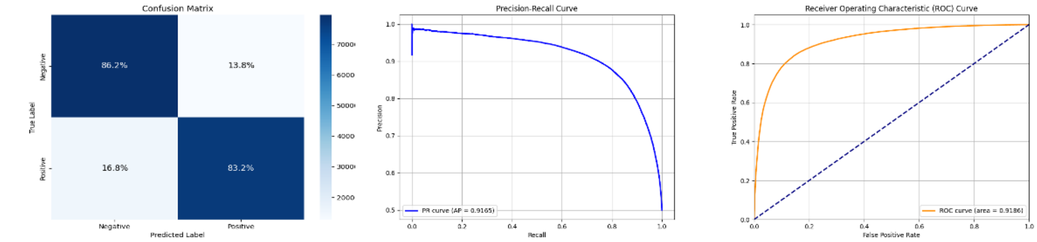
Confusion Matrix
True Label
Negative
Positive
86.2%
13.8%
16.8%
83.2%
Predicted Label
Precision-Recall Curve
Precision
Recall
PR curve (AP = 0.9165)
Receiver Operating Characteristic (ROC) Curve
True Positive Rate
False Positive Rate
ROC curve (area = 0.9186)

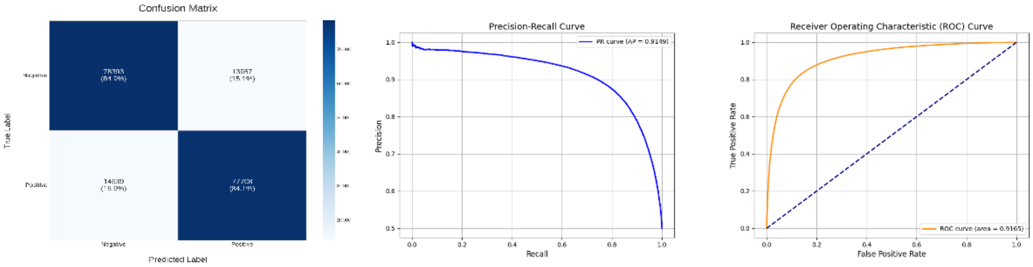
Confusion Matrix
True Label
Predicted Label
Precision-Recall Curve
Precision
Recall
PR curve (AP = 0.9149)
Receiver Operating Characteristic (ROC) Curve
True Positive Rate
False Positive Rate
ROC curve (area = 0.9165)

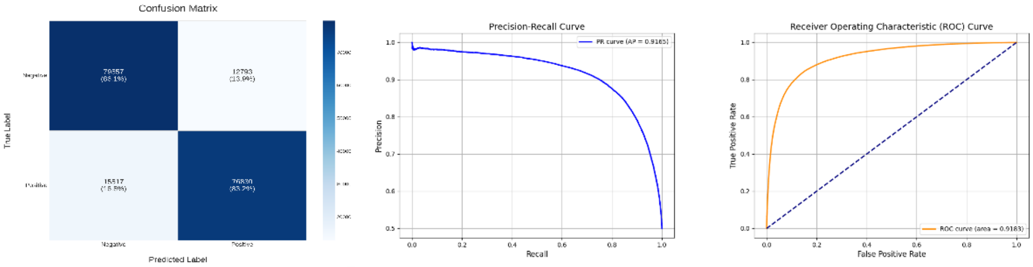
Confusion Matrix
True Label
Predicted Label
Precision-Recall Curve
Precision
Recall
PR curve (AP = 0.9165)
Receiver Operating Characteristic (ROC) Curve
True Positive Rate
False Positive Rate
ROC curve (area = 0.9183)

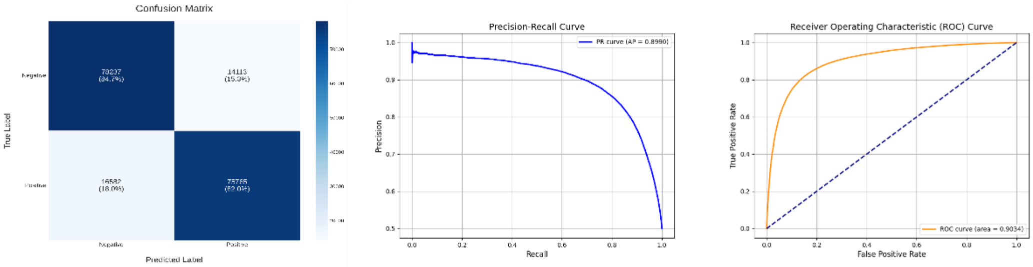
Confusion Matrix
True Label
Predicted Label
Precision-Recall Curve
Precision
Recall
PR curve (AP = 0.8990)
Receiver Operating Characteristic (ROC) Curve
True Positive Rate
False Positive Rate
ROC curve (area = 0.9034)

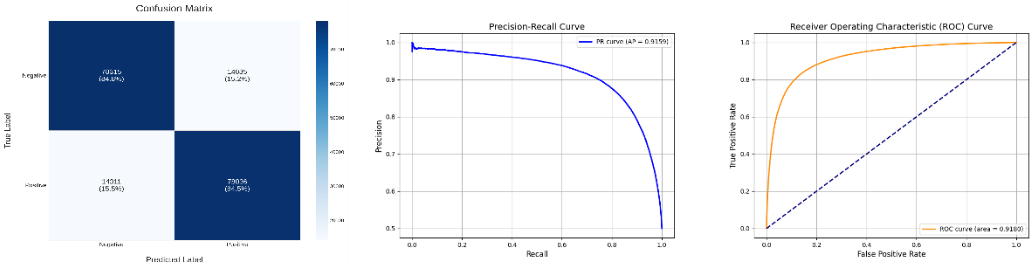
Confusion Matrix
True Label
Predicted Label
Precision-Recall Curve
Precision
Recall
PR curve (AP = 0.9159)
Receiver Operating Characteristic (ROC) Curve
True Positive Rate
False Positive Rate
ROC curve (area = 0.9180)

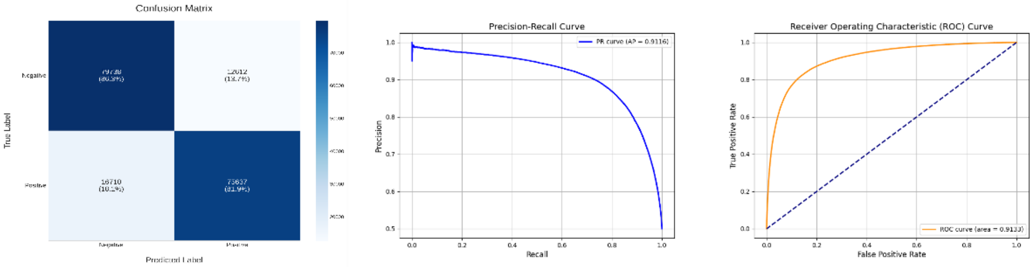
Confusion Matrix
True Label
Predicted Label
Precision-Recall Curve
Precision
Recall
PR curve (AP = 0.9116)
Receiver Operating Characteristic (ROC) Curve
True Positive Rate
False Positive Rate
ROC curve (area = 0.9133)

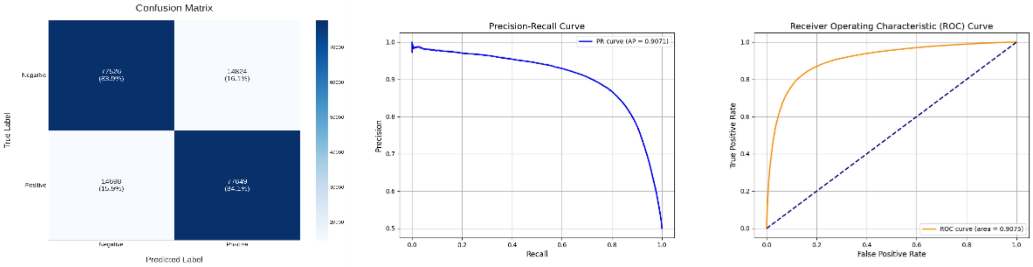
Confusion Matrix
True Label
Predicted Label
Precision-Recall Curve
Precision
Recall
PR curve (AP = 0.9071)
Receiver Operating Characteristic (ROC) Curve
True Positive Rate
False Positive Rate
ROC curve (area = 0.9075)

**Figure S3.** The performance of the model after removing each component was detailedly demonstrated. Confusion matrices, PR curves, and ROC curves were respectively plotted. The ablation operations from top to bottom were as follows: removing pre-training, removing LoRA, removing ESM2 feature extraction, removing the CNN module, removing the Transformer module, removing Cross-Multi-Attention feature fusion, and replacing Mimo Loss with Crossentropyloss.

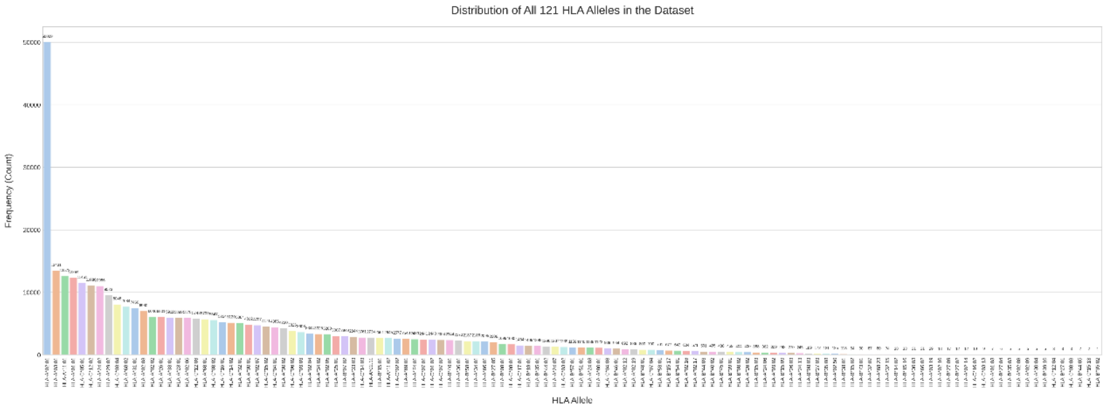


**Figure S4.** The frequency distribution of the PSGM dataset based on Allele classification. The x-axis represents the Allele type, and the y-axis represents the frequency statistics.

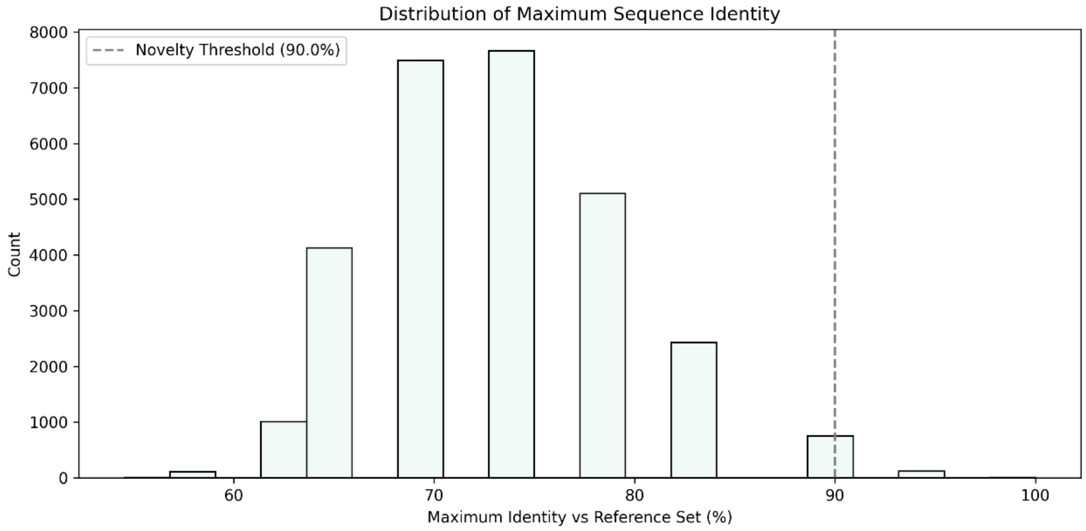


**Figure S5.** Maximum sequence identity distribution plot. Shows the frequency distribution of generated sequences based on their sequence identity to the training set (at a 90% threshold), with the majority of generated sequences falling within the 70-80% identity range.

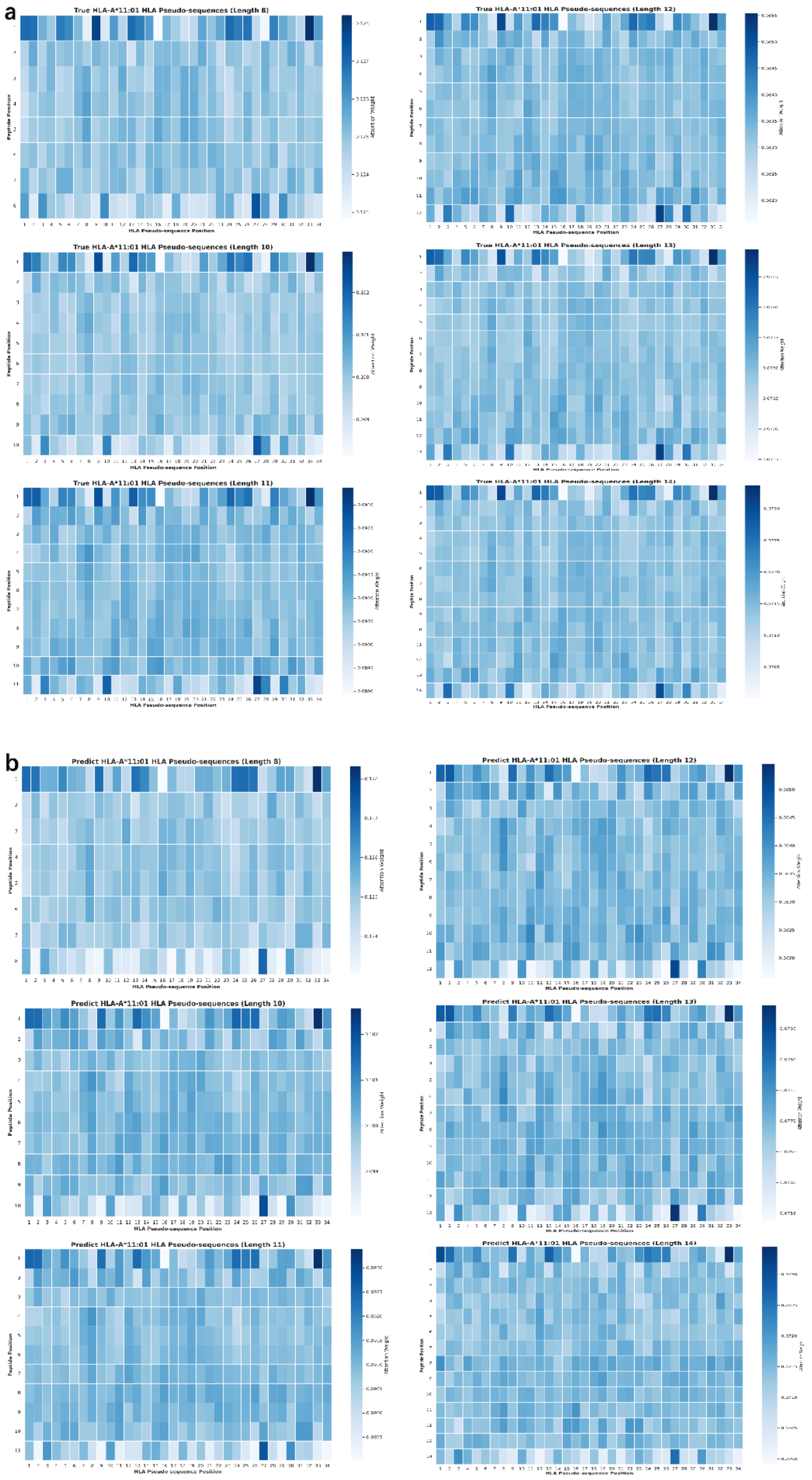


**Figure S6.** The figures respectively demonstrate the comparison of the attention distribution of the actual and predicted binding at various peptide segment lengths.

The horizontal axis represents the pseudo-sequence position, and the vertical axis represents the peptide segment position.

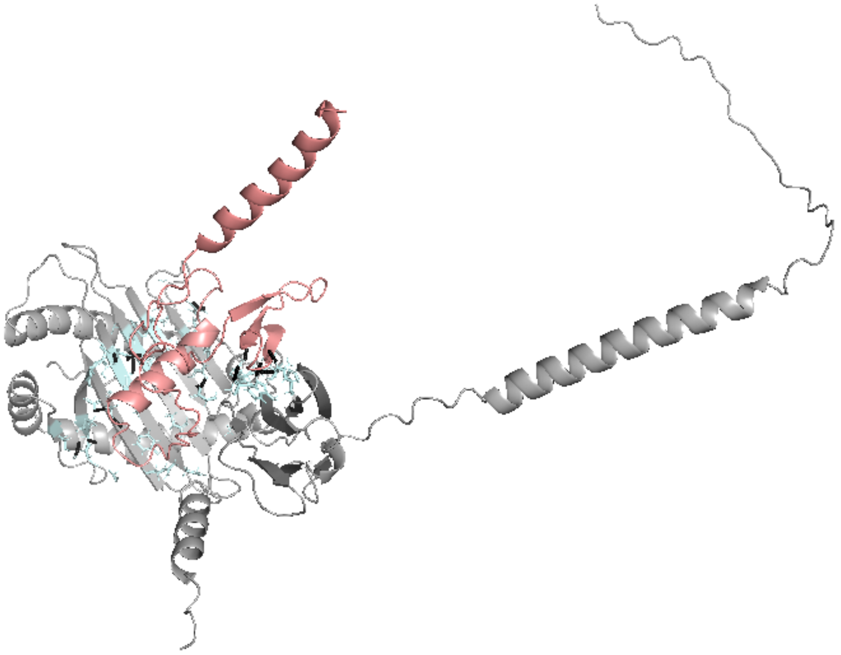

**Figure S7.** Structural modeling of a representative novel peptide-HLA-I complex identified by PSGM. A detailed view of the binding interface highlights the key interactions for peptide(ITDGQIFL) and HLA-A*11:01. Specific HLA-I residues that form the binding pocket are shown as cyan sticks. Hydrogen bonds, which indicate stable molecular interactions, are depicted as black dashed lines. The tight fit and the network of hydrogen bonds provide strong structural evidence for the biological plausibility of this novel interaction predicted by PSGM.

Table 1: Performance Metrics per Epoch for PHbinder without Pretrain

| Epoch | ACC | AUC | Loss | F1 | Recall | Precision | MCC |
|---|---|---|---|---|---|---|---|
| 1 | 83.58 | 90.67 | 0.876 | 83.25 | 81.49 | 85.09 | 0.6723 |
| 2 | 83.91 | 91.13 | 0.863 | 83.42 | 80.84 | 86.18 | 0.6796 |
| 3 | 84.21 | 91.32 | 0.860 | 84.15 | 83.67 | 84.63 | 0.6843 |
| 4 | 84.47 | 91.50 | 0.857 | 84.34 | 83.47 | 85.22 | 0.6896 |
| 5 | 84.39 | 91.53 | 0.855 | 84.32 | 83.77 | 84.87 | 0.6879 |
| 6 | 84.47 | 91.68 | 0.854 | 83.99 | 81.39 | 86.77 | 0.6907 |
| 7 | 84.28 | 91.68 | 0.853 | 83.54 | 79.66 | 87.82 | 0.6887 |
| 8 | 84.73 | 91.79 | 0.852 | 84.68 | 84.29 | 85.08 | 0.6947 |
| 9 | 84.52 | 91.68 | 0.851 | 84.10 | 81.79 | 86.56 | 0.6915 |
| 10 | 84.78 | 91.90 | 0.850 | 84.41 | 82.27 | 86.66 | 0.6965 |
| 11 | 84.74 | 91.87 | 0.849 | 84.43 | 82.62 | 86.32 | 0.6954 |
| 12 | 84.75 | 91.93 | 0.848 | 84.59 | 83.64 | 85.57 | 0.6951 |
| 13 | 84.58 | 91.94 | 0.847 | 84.81 | 85.97 | 83.68 | 0.6918 |
| 14 | 84.65 | 91.95 | 0.847 | 84.06 | 80.83 | 87.56 | 0.6952 |
| 15 | 84.90 | 92.00 | 0.846 | 84.72 | 83.64 | 85.83 | 0.6982 |
| 16 | 84.70 | 91.98 | 0.845 | 84.83 | 85.44 | 84.23 | 0.6940 |
| 17 | 84.89 | 92.00 | 0.844 | 84.74 | 83.83 | 85.67 | 0.6979 |
| 18 | 84.69 | 91.94 | 0.844 | 84.31 | 82.11 | 86.63 | 0.6949 |
| 19 | 84.65 | 91.92 | 0.843 | 84.77 | 85.27 | 84.27 | 0.6931 |
| 20 | 84.68 | 91.91 | 0.842 | 84.38 | 82.60 | 86.18 | 0.6941 |

Table 2: Performance Metrics per Epoch for PHbinder without Pretrain

| Epoch | ACC | AUC | Loss | F1 | Recall | Precision | MCC |
|---|---|---|---|---|---|---|---|
| 1 | 84.42 | 91.64 | 0.859 | 83.26 | 83.62 | 84.98 | 0.6981 |
| 2 | 85.32 | 92.23 | 0.850 | 85.11 | 83.79 | 86.47 | 0.7067 |
| 3 | 85.43 | 92.31 | 0.849 | 85.22 | 83.93 | 86.55 | 0.7089 |
| 4 | 85.38 | 92.32 | 0.848 | 85.16 | 83.81 | 86.56 | 0.7079 |
| 5 | 85.22 | 92.36 | 0.847 | 84.73 | 81.89 | 87.78 | 0.7061 |
| 6 | 85.44 | 92.37 | 0.845 | 85.45 | 85.40 | 85.51 | 0.7088 |
| 7 | 85.37 | 92.34 | 0.844 | 85.38 | 85.31 | 85.46 | 0.7075 |
| 8 | 85.29 | 92.32 | 0.843 | 84.90 | 82.60 | 87.33 | 0.7068 |
| 9 | 85.29 | 92.30 | 0.842 | 85.15 | 84.23 | 86.09 | 0.7059 |
| 10 | 85.27 | 92.30 | 0.840 | 85.07 | 83.77 | 86.40 | 0.7058 |
| 11 | 85.24 | 92.28 | 0.839 | 84.81 | 83.12 | 85.42 | 0.7055 |

Table 3: New Peptide-HLA-I Alleles

| Peptide | Best Allele | Mhcflurry Affinity |
|---|---|---|
| GSVYITLKK | HLA-A11:01 | 31.17 |
| EISSILKEL | HLA-A68:02 | 41.32 |
| ITDGQIFL | HLA-C05:01 | 30.02 |
| LLDVARTSL | HLA-C05:01 | 31.02 |
| HTSSAIPVPK | HLA-A34:02 | 27.63 |
| GTGASGSFK | HLA-A11:01 | 42.73 |
| EISSIISKM | HLA-A26:01 | 23.36 |
| ISVASFQEL | HLA-C03:03 | 34.14 |
| ASPSGSQL | HLA-C01:02 | 36.86 |
| IASAIVNEL | HLA-C03:03 | 24.06 |
| SAFLSAIFL | HLA-C17:01 | 37.89 |
| HADASSKVL | HLA-C05:01 | 26.46 |
| GMYVFLHAV | HLA-A02:11 | 11.78 |
| AQAPIAAVL | HLA-A02:05 | 33.36 |
| LAIPFAITI | HLA-C03:03 | 30.37 |
| TAKRSGAVF | HLA-C12:02 | 40.52 |
| VADGIFKAEL | HLA-C05:01 | 35.79 |
| VSHSTHRTF | HLA-C12:02 | 36.50 |
| YIDLPPPRL | HLA-C05:01 | 23.66 |
| SLIGNLHLA | HLA-A02:03 | 12.89 |
| LLDEVLNVM | HLA-C05:01 | 25.52 |
| FLDNLHINL | HLA-A02:07 | 18.09 |
| FVAPPTAAV | HLA-C03:03 | 25.85 |
| ALAKAAAAI | HLA-A02:02 | 17.56 |
| GTADVHFER | HLA-A68:01 | 25.99 |
| VAPVTHVSV | HLA-C01:02 | 31.11 |
| SFEQVVNELF | HLA-A29:02 | 313.44 |
| VTEIISEI | HLA-C05:01 | 87.81 |
| DLFFPGYSKGR | HLA-A34:01 | 110.11 |
| RVDFCGKGY | HLA-A29:02 | 168.78 |
| RQVVNVITTK | HLA-A03:01 | 68.28 |
| AYMELQQKAEF | HLA-C14:03 | 94.13 |
| NNALQNLAR | HLA-A68:01 | 205.25 |
| LPDGQVITI | HLA-C05:01 | 57.95 |

Table 4: Hyperparameter settings for the PHbinder model.

| Parameter | Value | Description |
|---|---|---|
| **I. General & Data Preprocessing** | | |
| Random Seed | 42 | Seed for all random number generators to ensure reproducibility. |
| Padding Length | 16 | All input peptide sequences were padded or truncated to this length. |
| Base Language Model | `esm2_t30_150M` | The ESM-2 model used, with features extracted from layer 30. |
| **II. LoRA Pre-training Phase** | | |
| Rank ($r$) | 16 | The rank of the LoRA update matrices. |
| Alpha ($\alpha$) | 32 | The LoRA scaling factor. |
| LoRA Dropout | 0.1 | Dropout rate for the LoRA layers. |
| Optimizer | AdamW | |
| Learning Rate | 1e-5 | |
| Batch Size | 32 | |
| Early Stopping | 5 | Patience for early stopping based on validation loss. |
| **III. PHbinder Main Model Architecture** | | |
| Rank ($r$) | 8 | The rank of the LoRA update matrices. |
| Alpha ($\alpha$) | 16 | The LoRA scaling factor. |
| LoRA Dropout | 0.1 | Dropout rate for the LoRA layers. |
| Transformer Layers | 6 | Number of layers in the Transformer encoder. |
| Attention Heads | 16 | Number of heads in the multi-head attention mechanism. |
| Model Dimension | 640 | Hidden dimension for Transformer and ESM-2. |
| Feed-Forward Dim | 64 | Inner dimension of the feed-forward network. |
| CNN Channels | 256 | Number of feature map channels in the CNN branch. |
| Classifier Dropout | 0.3 | Dropout rate before the final fully connected layer. |
| **IV. PHbinder Main Training Phase** | | |
| Optimizer | Adam | |
| Learning Rate | 5e-6 | |
| Weight Decay | 0.0025 | |
| Batch Size | 64 | |
| Early Stopping | 5 | Patience based on validation accuracy. |
| Loss Function | Mimoloss | |

Table 5: Hyperparameter settings for the PSGM model.

| Parameter | Value | Description |
|---|---|---|
| **I. General & Data Configuration** | | |
| Peptide Max Length | 14 | Maximum length for input peptide sequences. |
| HLA Pseudo-seq Length | 34 | Fixed length of the target and generated HLA pseudo-sequences. |
| Conditional Encoder | `esm2_t30_150M` | Pre-trained language model for encoding input peptides. |
| ESM-2 Feature Layer | 30 | Peptide representations are extracted from the 30th layer. |
| **II. Generator Architecture** | | |
| Embed Dimension | 256 | Internal working dimension for model components. |
| Decoder Layers | 6 | Number of Transformer layers in the decoder. |
| Attention Heads | 8 | Number of heads in the multi-head attention. |
| Feed-Forward Dim | 1024 | Inner dimension of the feed-forward networks in the decoder. |
| **III. Discriminator Architecture** | | |
| Embed Dimension | 256 | Input embedding dimension for the discriminator. |
| Encoder Layers | 3 | Number of Transformer layers in the encoder. |
| **IV. GAN Training & Optimization** | | |
| Batch Size | 128 | |
| Optimizer | AdamW | Used for both Generator and Discriminator. |
| Learning Rate | 1e-7 | |
| Adversarial Weight ($\lambda_{\text{adv}}$) | 0.3 | Weight to balance generative and adversarial objectives. |
| Gradient Clipping | 1.0 | Maximum norm for the generator's gradients. |
| Early Stopping | 3 | Patience for early stopping based on validation loss. |
| **V. Inference & Constrained Generation** | | |
| Decoding Strategy | Top-p Sampling | Strategy to sample the next amino acid from the probability distribution. |
| Top-p ($p$) | 0.9 | Cumulative probability threshold for nucleus sampling. |
| Temperature ($\tau$) | 1.0 | Controls the smoothness of the probability distribution. |
| Constrained Decoding | Position Masks | Ensures each position is sampled from a predefined set of valid amino acids. |
